# Rec8 cohesin-mediated axis-loop chromatin architecture is required for meiotic recombination

**DOI:** 10.1101/2021.12.09.472021

**Authors:** Takeshi Sakuno, Sanki Tashiro, Hideki Tanizawa, Osamu Iwasaki, Da-Qiao Ding, Tokuko Haraguchi, Ken-ichi Noma, Yasushi Hiraoka

**Affiliations:** Graduate School of Frontier Biosciences, Osaka University, Suita 565-0871, Japan; Institute of Molecular Biology, University of Oregon, Eugene, OR 97403, USA; Advanced ICT Research Institute Kobe, National Institute of Information and Communications Technology, Kobe 651-2492, Japan; Institute for Genetic Medicine, Hokkaido University, Sapporo 060-0815, Japan

## Abstract

During meiotic prophase, cohesin-dependent axial structures are formed in the synaptonemal complex (SC). However, the functional correlation between these structures and cohesion remains elusive. Here, we examined the formation of cohesin-dependent axial structures in the fission yeast *Schizosaccharomyces pombe*. This organism forms atypical SCs composed of linear elements (LinEs) resembling the lateral elements of SC but lacking the transverse filaments. Hi-C analysis using a highly synchronous population of meiotic *S. pombe* cells revealed that the axis-loop chromatin structure formed in meiotic prophase was dependent on the Rec8 cohesin complex. In contrast, the Rec8-mediated formation of the axis-loop structure occurred in cells lacking components of LinEs. To dissect the functions of Rec8, we identified a *rec8-F204S* mutant that lost the ability to assemble the axis-loop structure without losing cohesion of sister chromatids. This mutant showed defects in the formation of the axis-loop structure and LinE assembly and thus exhibited reduced meiotic recombination. Collectively, our results demonstrate that the Rec8-dependent axis-loop structure provides a structural platform essential for LinE assembly, facilitating meiotic recombination of homologous chromosomes, independently of its role in sister chromatid cohesion.

## INTRODUCTION

Meiosis is a process of cell division unique to sexual reproduction, which halves the chromosome number in two successive nuclear divisions after a single round of DNA replication to produce haploid gametes. In the first meiotic division (meiosis I), homologous chromosomes (homologs) are segregated to opposite poles. This type of division is called reductional segregation and is essentially different from mitosis or meiosis II, in which sister chromatids are segregated to opposite poles (1). Crossovers formed by reciprocal recombination constitute a physical link between homologs, ensuring reductional segregation (2,3). Failure to establish such interhomolog links leads to the mis-segregation of chromosomes during meiosis I, resulting in aneuploidy of gametes. This failure is a major cause of human miscarriage or developmental disorders caused by chromosome trisomy, such as Down syndrome (4,5).

During meiotic prophase, homologs become closely associated or aligned through an elaborate proteinaceous structure called the synaptonemal complex (SC), followed by crossover recombination. The SC consists of two lateral elements connected by a central region (6,7). The lateral element is formed along the axis joining a pair of sister chromatids and is essential for meiotic recombination between homologs. Formation of these meiosis-specific chromosome structures is mediated by the meiotic cohesin complex (1,8).

The cohesin complex consists of four protein components: two structural maintenance of chromosome (SMC) components, one kleisin component, and one stromal antigen (SA) component (9-12); these components are called Smc1α, Smc3, Rad21/Scc1, and SA1 or SA2, respectively, in human somatic cells (Rad21 cohesin complex, hereafter). The somatic Rad21 cohesin complex establishes cohesion of a pair of replicated sister chromatids. Replacement of kleisin during meiosis occurs commonly in most organisms, in which the somatic kleisin Rad21/Scc1 is replaced by Rec8 (Rec8 cohesin complex, hereafter) (9-12). In addition to sister chromatid cohesion, the meiotic Rec8 cohesin complex is required for the assembly of lateral elements in diverse eukaryotes (8,13-22). A recent study using high-throughput chromosome conformation capture (Hi-C) (23) in the budding yeast *Saccharomyces cerevisiae* clearly revealed that the Rec8 cohesin complex is essential to construct meiosis-specific chromatin loop structures (24). As studied in somatic cells, cohesin complexes can form higher-order structures by pulling together distant regions of the chromosome, for which a loop extrusion model has been proposed (25-30). Pds5 and Wpl1/Wapl are cohesin-associating factors conserved in yeasts and vertebrates. These factors are considered to dissociate cohesin from chromatin (31,32), thus affecting higher-order chromosome structures (33-41).

The fission yeast *Schizosaccharomyces pombe* forms an atypical SC composed of a structure called the linear element (LinE). The LinE resembles the lateral element of the SC but lacks a central region composed of transverse filaments (42-47). The assembly of *S. pombe* LinEs requires phosphorylated Rec11, a homolog of the meiosis-specific SA3 subunit of the Rec8 cohesin complex; phosphorylated Rec11 interacts with Rec10, a homolog of the lateral element factor SYCP2 in mammals (*S. cerevisiae* Red1) (48,49). In LinE-defective mutants of *S. pombe* meiosis, Rec8 cohesin-dependent chromosomal structure is maintained (39,48). Similarly, in SYCP2-depleted mouse spermatocytes defective in lateral element formation, normal cohesin-axis is formed (22,50,51). Therefore, the Rec8-dependent chromosomal structure is independent of lateral elements (or LinEs in *S. pombe*), but the biological significance of this structure remains unclear.

In this study, we performed Hi-C analysis to delineate the meiotic chromosome structures of *S. pombe*. We then sought to clarify the functional significance of the Rec8-dependent axis-loop structures by identifying *rec8* mutants that specifically show defects in axis formation but not in cohesion.

## MATERIALS AND METHODS

### Schizosaccharomyces pombe strains

All media and growth conditions were as described previously (52). Complete medium supplemented with uracil, lysine, adenine, histidine, and leucine (YE5S), minimal medium (EMM2), and sporulation-inducing medium (SPA and SSA) were used unless otherwise stated. The strains used in this study are listed in Table S1. Deletion of the endogenous genes (*rec12*^*+*^, *rec11*^*+*^, *rec7*^*+*^, *rec15*^*+*^, *mek1*^*+*^, *rec8*^*+*^, *rec10*^*+*^, *hop1*^*+*^, *wpl1*^*+*^, *pds5*^*+*^, and *mei4*^*+*^) and tagging (*rec10*^*+*^, *rec8*^*+*^, *psm1*^*+*^, *rec25*^*+*^, *rad21*^*+*^, and *rec11*^*+*^) with GFP, mCherry, 3×hemagglutinin (HA) tag, 3×Pk (V5), or 3×Flag were performed according to the PCR-based gene targeting method for *S. pombe*, using the *ura4*^*+*^, *kanMX6* (*kanR*), *hphMX6* (*hygR*), *natMX6* (*natR*), and *bsdR* genes as selection markers (53-55). Histone H2B tagging by mCherry for time-lapse imaging has been described previously (56). The CK1 shutoff mutant, *rec11-5A* mutant, *rec11-5D* mutant, and the strain expressing Rec10-GFP-Rec11 fusion protein have been described previously (48). For the assay of crossing over recombination frequency, the *hygR* cassette was inserted 1,215 bp upstream from the start codon of the *cox7*^*+*^ gene, and the *natR* cassette was inserted 1,000 bp upstream from the start codon of *vps38*^*+*^ gene, using a PCR-based gene targeting method. To generate strains of *rec8* point mutants with tagging of GFP or 3×Pk tag, the endogenous *rec8*^*+*^ locus with its promoter and terminator was cloned into pUC119. Using the NEBuilder system (New England Biolabs), a DNA fragment encoding GFP or 3×Pk tag with *kanR or bsdR* was introduced immediately before the stop codon of the *rec8* gene cloned in pUC119. Next, target sequences of *rec8*^*+*^ were subsequently mutated using the KOD One Master Mix Blue (TOYOBO) to generate an amino acid substitution. In case of *rec8-S552P*, a C-terminal portion of the *rec8*^*+*^ open reading frame, including the mutation site and marker gene with the *rec8*^*+*^ terminator, was amplified by PCR and transformed into the wild-type *rec8*^*+*^ locus. Alternatively, in the case of N-terminal mutations such as *rec8-F204S*, the *rec8*^*+*^ gene region spanning the promoter to the terminator, including the mutation and marker gene, was amplified by PCR and transformed into the Δ*rec8::ura4*^*+*^ strain. Correct replacement was confirmed using PCR and direct sequencing. The strain in which the mitotic cohesins, Rad21 and Psc3, were exchanged for meiotic cohesins (Rec8 and Rec11) has been described previously (57). To construct the *pat1-as2* diploid with MDR-sup, haploid strains (*h*^*-*^ *pat1-as2 MDR-sup* and *h*^*-*^ *pat1-as2 MDR-sup mat-Pc*) were used (they were generously provided by Masamitsu Sato, Waseda University). After introducing a tag or disrupting the gene of interest in both haploids, protoplast fusion was performed as described previously (58), except for the use of Lysing Enzyme (Sigma-Aldrich) instead of Lallzyme or Zymolyase.

### Preparation of meiotic cells

For microscopic observation of Rec8-GFP, Rec11-GFP, Rec10-mCherry, and Rec25-mCherry, *h*^*90*^ cells cultured on YE plates at 26°C overnight were streaked on SSA plates or suspended in 20 mg/mL of leucine and spotted on SPA or SSA plates. The cells were incubated at 26°C for 6–10 h to observe the cells at the horsetail stage. In case of *lacO*/lacI-GFP observation for cohesion assays (Figure 5C and S5C), *h*^*+*^ and *h*^*-*^ cells were mixed with 20 mg/mL of leucine and spotted on SSA or SPA plates. For the immunoprecipitation of Rec8-GFP with Psm1-3HA (Figures S5E), *h*^*+*^/*h*^*-*^ diploid cells with *mei4Δ* were used. The diploid cells were grown in EMM2 liquid medium containing 5 mg/mL NH_4_Cl at a density of 5 × 10^6^ cells/mL at 25°C and then resuspended in EMM2 (1% glucose) medium lacking NH_4_Cl at a density of 1 × 10^7^ cells/mL at 25°C for 15 h. For the immunoprecipitation of Rec8-3×Flag with Psm1 (Figure S5F), ChIP assay (Figure S6B), detection of phosphorylated Rec11 (Figure 7C), and Hi-C analyses, we used *h*^*-*^/*h*^*-*^ diploid cells with *pat1-as2* and *MDR-sup* alleles. Cells grown in YE liquid medium at a density of 5–10 × 10^6^ cells/mL at 25°C were collected by centrifugation and washed twice with EMM2 (1% glucose) medium lacking NH_4_Cl (EMM2-N). Cells were then resuspended in EMM2-N with 50 mg/mL leucine at a density of 5 × 10^6^ cells/mL and incubated at 25°C for 6.5 h to induce G1 arrest. By adding the same volume of EMM2-N with 50 mg/mL leucine and 15 µM 1-NM-PP1 (Toronto Research Chemicals Inc.) (5–8 × 10^6^ cells/mL), cells can enter meiosis. The culture was further incubated for 2.5 h or more. Before harvesting the meiotic cells, 1 mM phenylmethylsulfonyl fluoride (PMSF; Sigma-Aldrich) was added to reduce protein degradation.

### Rec8 mutant screening

The *rec8*^*+*^ cDNA mutant library used for the screening was constructed as follows. The DNA region encoding *3×HA-ura4*^*+*^of the plasmid VP068 (pUC119 base, *Padh1-rec8*^*+*^cDNA*-3×HA-ura4*^*+*^*-Trec8*) (59) was replaced by the DNA region encoding GFP-BsdR to produce VP068-GFP-bsd. Error-prone PCR of *rec8*^*+*^ cDNA in VP068-GFP-BsdR by Ex Taq DNA Polymerase (TaKaRa Bio) was performed using several concentrations of MnCl_2_ (final 0.05 mM, 0.15 mM, 0.25 mM, and 0.3 mM) and an additional 1 mM dGTP or dATP with 0.2 mM of dNTP mix (total eight conditions). By using the NEBuilder system, amplified *rec8*^*+*^cDNA pools (from eight conditions) were then cloned into the fragment, which was amplified by inverse PCR of VP068-GFP-BsdR without the *rec8*^*+*^ cDNA region. In total, approximately 5,300 *Escherichia coli* transformants (representing library size) were generated. All colonies were recovered from the plates, and plasmids (pRec8 library) were extracted. The pRec8 library was estimated to contain approximately six mutations per kilobase. Using the pRec8 library as a template, the region spanning 620 bp upstream of the *adh1*^+^ promoter to 600 bp downstream of the *rec8*^*+*^ terminator was amplified by PCR. These fragments were used to transform the host strains for screening. More than 95% of blasticidin-resistant colonies obtained after transformation showed correct integration, as confirmed by PCR. Rec8-GFP signals during meiotic prophase were observed among the blasticidin-resistant colonies, which showed relatively normal growth.

### Fluorescence microscopy

Fluorescence images were obtained using an Axioplan2 microscope (Carl Zeiss) equipped with a Quantix cooled CCD camera (Photometrics) and a DeltaVision microscope system (GE Healthcare Inc.) equipped with a CoolSNAP HQ^2^ cooled CCD camera (Photometrics) and a 60× Plan-ApoN SC oil immersion objective lens (NA = 1.40; Olympus). The brightness of the images was adjusted using Fiji software (60) without changing the gamma settings. The pairing frequency was quantified as described previously (61). For time-lapse live-cell imaging of Rec8-GFP and H2B-mCherry, cells were cultured overnight in liquid EMM2 at 26°C to reach the stationary phase; thereafter, the cells were spotted onto SPA plates to induce meiosis, followed by incubation at 26°C for 4–5 h. Zygotes suspended in 1% glucose EMM2 (1% glucose) without nitrogen were sonicated briefly and immobilized on a coverslip coated with lectin in a 35 mm glass-bottomed culture dish (MatTek) and observed at 26°C in liquid EMM2 (1% glucose) without nitrogen using a DeltaVision microscope in a temperature-controlled room. Time-lapse images were collected every 5 min at 9 focal sections with 0.45 μm interval at each time point. Projected images were subsequently acquired using Z-sectioning and projected according to the SoftWoRx software program (Applied Precision).

### Immunoprecipitation

The cell extracts (∼8 × 10^7^ cells) were prepared by bead-beating cells in IP buffer [50 mM HEPES-KOH at pH 8.0, 150 mM NaCl, 1 mM dithiothreitol, 2.5 mM MgCl_2_, 20% glycerol, 1 mM PMSF, 0.8% Triton X-100, 0.1% sodium deoxycholate, and 5× protease inhibitor cocktail (P8215; Sigma-Aldrich)]. Fifty units of DNase I (TaKaRa Bio) were added to the cell extract, and sonication was conducted (Branson Inc.). After collecting the supernatant as whole-cell extracts by centrifugation at 15,000 rpm for 30 min, immunoprecipitation was performed by incubating these extracts for 1 h at 4°C with anti-GFP polyclonal antibodies (Rockland) or anti-Flag polyclonal antibodies (anti-DDDDK, MBL) and an additional 2 h at 4°C with 20 μL of a slurry of IgG-conjugated Dynabeads (Invitrogen). After washing the beads five times with IP buffer, the whole-cell extracts and immunoprecipitates were subjected to western blotting using monoclonal anti-GFP (JL-8, Roche), Flag-M2 monoclonal antibody (Sigma-Aldrich), monoclonal anti-HA (3F10, Roche), and monoclonal anti-tubulin (DM1A, Abcam) antibodies.

### ChIP assay

The procedures were carried out as described previously (62). Anti-FLAG M2 monoclonal antibody (Sigma-Aldrich) and IgG-conjugated Dynabeads (Invitrogen) were used for immunoprecipitation. DNA prepared from the whole-cell extracts or immunoprecipitated fractions was analyzed via quantitative PCR (qPCR) with the StepOnePlus Real-Time PCR system (Thermo Fisher Scientific) using Power SYBR Green PCR Master Mix^©^ Green I Master (Thermo Fisher Scientific). The primers used for PCR have been described previously (48). Immunoprecipitation with control mouse IgG antibody (Abcam) was performed in each experiment to account for nonspecific binding in the ChIP fractions and verify the significance of immunoprecipitation. The percent IP was quantified using three independent qPCR experiments.

### Recombination assay

For intergenic recombination, haploid strains containing appropriate markers were mated and sporulated at 26°C on SSA plates for 30 h. The spores were isolated using β-glucuronidase solution (Wako) and plated on YE. The colonies were then replicated in selective media [EMM2-uracil, EMM2-lysine, EMM2-lysine, YE5S containing hygromycin B (FUJIFILM) or YE5S containing clonNAT (WERNER BioAgents)] and YE5S containing phloxine B (Wako) to select haploid cells. In the *ura1-lys3* interval, the degree of lysine auxotrophy among the uracil prototrophic colonies (n >700, replicated three times) was assessed. In the *cox7*^*+*^*-vps38*^*+*^ interval, the degree of hygromycin B-resistant (*hygR*-containing) colonies among the clonNAT-resistant (*natR*-containing) colonies was assessed (n >500, and replicated three times). In the *vps38*^*+*^*-leu1*^*+*^ interval, the degree of leucine prototrophy among the clonNAT-resistant (*natR*-containing) colonies was assessed (n >500, and replicated three times). For intragenic recombination, *h*^*90*^ haploid strains containing the *ade6-M26* allele and plasmids harboring the *ade6-469* allele according to the *ura4*^*+*^ marker (63) were sporulated at 26°C on SPA plates for 30 h. The spores were isolated by treatment with glusulase and plated on YE5S containing phloxine B (to count all viable haploid spores) and EMM2-adenine plates. The frequency of intragenic recombination was quantified as the degree of adenine prototrophic colonies among viable haploid spores (n >7,400, replicated three times).

### Flow cytometry

A total of 1 × 10^7^ cells were fixed with 70% ethanol and incubated with 50 μg/mL RNase A in 500 μL of 50 mM sodium citrate (pH 7.5) for 4 h at 37°C and then stained with 0.5 μg/mL propidium iodide. DNA content of 1.5 × 10^5^ cells was measured on a FACS Calibur instrument (BD Biosciences).

### In situ Hi-C

In situ Hi-C experiments were performed as described previously (64) with minor modifications. Asynchronous or meiotic cell cultures of 4 × 10^8^ cells were fixed with 3% paraformaldehyde at 26°C for 10 min. The fixed cells were disrupted using glass beads and Mini-Beadbeater-16 (BioSpec Products) in lysis buffer [50 mM HEPES/KOH pH 7.5, 140 mM NaCl, 1 mM EDTA, 1% Triton X-100, 0.1% sodium deoxycholate, 1× Complete (Roche, 11836170001), and 1 mM PMSF]. The lysed cells were collected by centrifugation; pellets were resuspended with 1.11× NEBuffer 2 (B7002, NEB) containing 0.1% sodium dodecyl sulfate (SDS), and cells were permeabilized by incubation at 62°C for 7 min. After SDS was quenched by adding 1/9 volume of 10% Triton X-100, the permeabilized cells were treated with 25 units of MboI (R0147, NEB) at 37°C overnight. The enzyme was inactivated by incubation at 62°C for 20 min. After centrifugation, the pellet containing the restricted DNA fragments was used in the following procedure. A half volume of fill-in mix [1× NEBuffer 2, 150 µM biotin-14-dATP (NU-835-BIO14, Jena Bioscience), 150 µM dCTP, 150 µM dGTP, 150 µM dTTP, 0.4 units/µL Klenow fragment (M0210, NEB)] was added to the pellet and incubated at 37°C for 45 min. Subsequently, three volumes of ligation mix [1.33× T4 DNA ligase buffer (B0202, NEB), 1.11% Triton X-100, 133 µg/mL bovine serum albumin (B9001, NEB), 5.56 units/µL T4 DNA ligase (M0202, NEB)] was added, followed by incubation at room temperature (∼25°C) for 4 h. After ligation, the pellet was collected by centrifugation, resuspended in proteinase K solution [10 mM Tris-HCl pH 8.0, 200 mM NaCl, 1 mM EDTA, 0.5% SDS, 0.5 mg/mL proteinase K (25530049, Invitrogen)], and incubated at 65°C overnight. DNA was purified by phenol-chloroform treatment followed by ethanol precipitation and then fragmented by sonication with a model UCD-200 Bioruptor (Diagenode). After the biotin-labeled DNA was pulled down with Dynabeads Streptavidin T1 (65601, Invitrogen), the beads were washed twice with TNB (5 mM Tris-HCl pH 7.5, 0.5 mM EDTA, 1 M NaCl, 0.05% Tween-20) at 55°C for 2 min. To ligate the Illumina sequencing adaptors to the biotinylated DNA fragments, the beads were successively treated with end repair mix [1× T4 DNA ligase buffer, 1 mM dATP, 0.5 mM each dNTP, 0.5 units/µL T4 polynucleotide kinase (M0201, NEB), 0.12 units/µL T4 DNA polymerase (M0203, NEB), 0.05 units/µL Klenow fragment] at 25°C for 30 min, dA-tailing mix [1× NEBuffer 2, 0.5 mM dATP, 0.25 units/µL Klenow exo-(M0212, NEB)] at 37°C for 30 min, and Illumina adaptor ligation mix [0.075x End repair reaction buffer (E6050, NEB), 0.32× Ligation master mix (E7595, NEB), 0.011× Ligation enhancer, and 45 nM NEBNext adaptor for Illumina] at 20°C for 15 min followed by 0.032 units/µL USER treatment at 37°C for 15 min, with two washes with TNB at 55°C for 2 min after each reaction. The resultant adaptor-ligated Hi-C library was amplified by PCR with five to seven cycles with Q5 master mix (M0544, NEB) and index primers (E7335/E7500/E7710/E7730, NEB) before 37 bp paired-end sequencing using the NextSeq 500 device (Illumina).

### In situ Hi-C data analysis

In situ Hi-C data were processed as described previously (64). The mapping statistics of Hi-C data are summarized in Table S2. The paired reads were separately aligned to the fission yeast genome (version ASM294v2.19) using Bowtie2 (version 2.2.9) with the iterative alignment strategy. The aligned reads were assigned to MboI fragments. Redundant paired reads derived from PCR biases, aligned to repetitive sequences or with low mapping quality (MapQ < 30) were removed. Paired reads potentially derived from self-ligation and undigested products were also discarded. The fission yeast genome was divided into non-overlapped 20, 5, or 2 kb bins. Raw contact matrices were constructed by counting paired reads assigned to two bins, and Hi-C biases in the contact matrices were corrected using the ICE method (65). The ICE normalization was repeated 30 times. ICE-normalized contact scores were used for all heat maps, where the maximum intensity corresponded to the top 5% score. For subtraction heat maps, the top 5% absolute values of difference scores (*T*) were extracted. The *T* and **-***T* scores were drawn as the maximum and minimum intensity, respectively.

### Loop detection

Knight and Ruiz (KR) normalized matrices (66) in hic format were calculated from raw contact matrices using the “pre” function of Juicer_tools (version 1.22.01) (67). Loop calling was performed for KR-normalized matrices using HiCCUPS (“hiccups” function of Juicer_tools) with the following parameters: r 2000 -k KR -f .01 -p 3 -i 7 -d 10000”.

## RESULTS

### Induction of highly synchronized meiosis under physiological conditions

To perform Hi-C analysis of chromosomal configurations during meiotic prophase, we first developed an experimental system to prepare *S. pombe* cell populations undergoing highly synchronized meiotic progression under physiological conditions. For this purpose, we employed an *h*^*-*^*/h*^*-*^ *pat1-as2/pat1-as2* diploid strain harboring one copy of the *mat*-Pc gene cassette (68-70). In this strain harboring an ATP analog-sensitive allele of the Pat1 kinase (*pat1-as2*), meiosis can be induced by addition of the ATP analog that inactivates Pat1 kinase. This allele offers advantages over the temperature-sensitive allele (*pat1-114*), in which raising the temperature causes detrimental effects on meiosis (68,71). In addition, MDR-sup alleles (72,73), which allow efficient cellular uptake of ATP analogs, were also exploited. The meiotic progression of wild-type diploid cells harboring Rec8-GFP was highly synchronous (∼70%), as indicated by the timing of nuclear division in meiosis I (Figure S1A). Pre-meiotic DNA replication was almost completed approximately 1.5 h after the induction of meiosis by the addition of an ATP analog after G1 arrest via nitrogen starvation (Figure S1A, left panel).

### Hi-C analysis recapitulates meiotic bouquet chromosomes in *S. pombe*

For Hi-C analysis, cells were collected at 0 h (G1) and 2.5 h (meiotic prophase) after meiosis induction. Cells in vegetative growth (VG; mainly in mitotic G2 phase) were also collected for comparison. Notably, even at 0 h, a significant amount of Rec8-GFP protein was expressed and loaded onto chromatin (Figures S1B and S1C). The amount of Rec8-GFP peaked at approximately 2.5–3 h after meiotic induction (Figures S1B and S1C). We detected obvious centromere clustering (indicated by green arrows in Figures 1B and 1C) in the Hi-C contacts of all three chromosomes in VG cells, suggesting a Rabl orientation (Figure 1A, left). In contrast, as cells progressed from G1 to meiotic prophase, centromere clustering became vague, while telomere clustering became evident (Figures 1B and 1C; blue arrows). As cells entered the meiotic prophase, X-shaped inter- and intra-chromosomal contacts appeared (Figures 1B and 1C). These X-shaped contact patterns represent chromosomal arms bundled at the telomere, as previously demonstrated in *S. pombe* meiotic prophase (74,75) (Figure 1A; right). In *S. pombe*, alignment of homologs relies on oscillatory movement of the nucleus (horsetail movement) in a bouquet orientation with all telomeres clustered to the spindle pole body (SPB) (42,75-79). To systematically evaluate the degree of alignment of intra-chromosomal arms, we devised a measure “alignment index” (Figure S1D). The alignment index was calculated as the shortest distance from the telomere that gives normalized Hi-C contact scores above the threshold (refer to the legend of Figure S1D for details). It corresponds to a length of the solid vertical line in Figure 1D. The alignment index of chromosome 1 increased from 1.6 (onset of meiosis at 0 h) to 2.5 (meiotic prophase at 2.5 h) compared with 0.55 in VG (Figure 1D). A similar structure was observed during meiotic prophase in mouse spermatocytes by Hi-C analysis, although it was not as evident as in *S. pombe* (80).

**Figure 1.**
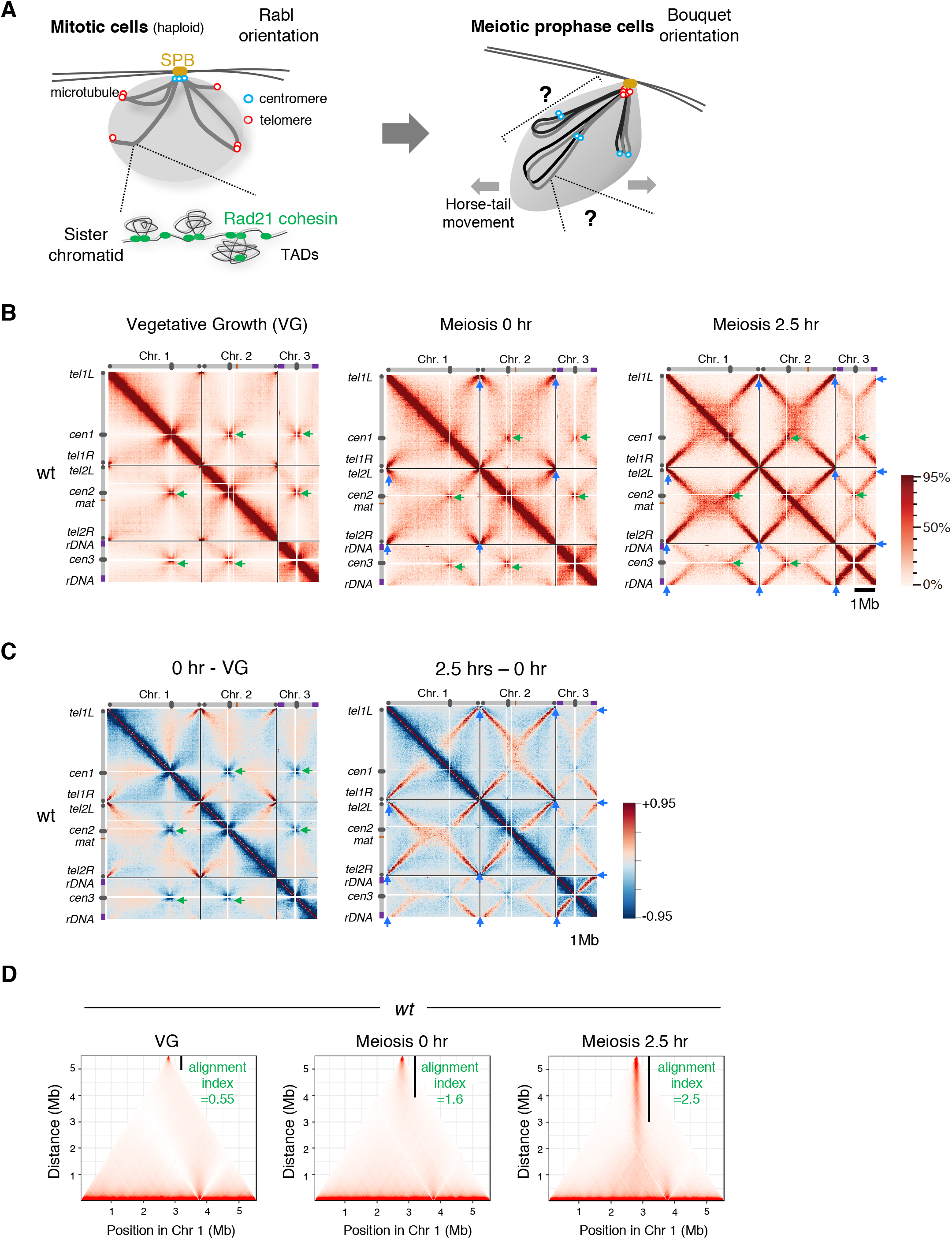
Hi-C analysis firmly recapitulates the structure of meiotic bouquet chromosomes. (A) Chromosome arrangement in mitosis and meiosis in *Schizosaccharomyces pombe*. In mitosis (left), centromeres are clustered at the spindle pole body (SPB) and display the Rabl orientation. The grey lines indicate sister chromatids. In meiotic prophase (right), grey and black lines indicate a homologous pair of chromosomes. Chromosomes are bundled by the telomeres clustered at the SPB, representing the bouquet orientation. (B) Hi-C contact maps at 20 kb resolution for three chromosomes prepared at vegetative growth (VG) and 0 and 2.5 h after entry into meiosis. Schematic views of each chromosome are shown on the top and the left. Green arrowheads represent the centromere clustering. Blue arrowheads in meiosis 0 and 2.5 h represent the telomere clustering as depicted by the left lower diagonal lines. The color scale on the right represents the percent ranking of contact scores. (C) Heat maps showing the difference of contacts detected by Hi-C. Left, subtraction of VG data from 0 h data. Right, subtraction of 0 h data from 2.5 h data. The color scale on the right represents the percent ranking of difference scores. (D) Degree of alignment of the two chromosome arms of telomere-bundled chromosome 1 is quantified as the alignment index (Figure S1D) and is shown as the value along with a black vertical line in the Hi-C contact map.

### Rec8 cohesin complex produces a chromosome shape suitable for alignment

To examine contributions of LinE and meiotic recombination to the chromosome structures, we performed Hi-C analysis in mutant cells lacking three factors that play a central role in meiotic recombination (Rec10, Rec12, and Rec8). Rec10 (a homolog of Red1/SYCP2) is a major component of LinE, and Rec12 is a Spo11 homolog that induces DNA double-strand breaks (81,82). The Rec8 cohesin complex is required for proper assembly of LinE (46,48,49,83). For Hi-C analysis, we first confirmed synchronous progression of meiosis in these mutants (Figure S2). Hi-C analysis for *rec10Δ* and *rec12Δ* cells showed X-shaped contacts similar to wild-type cells (Figures 2A and 2B), with an alignment index of 2.8 (*rec10Δ*), 2.5 (*rec12Δ*), and 2.5 (*wild-type, wt*). In contrast, *rec8Δ* cells showed prominent defects with almost no X-shaped contacts (Figures 2A and 2B; alignment index 1.5). These results suggest that Rec8 is required for the alignment of homologs in a manner distinct from LinE formation.

**Figure 2.**
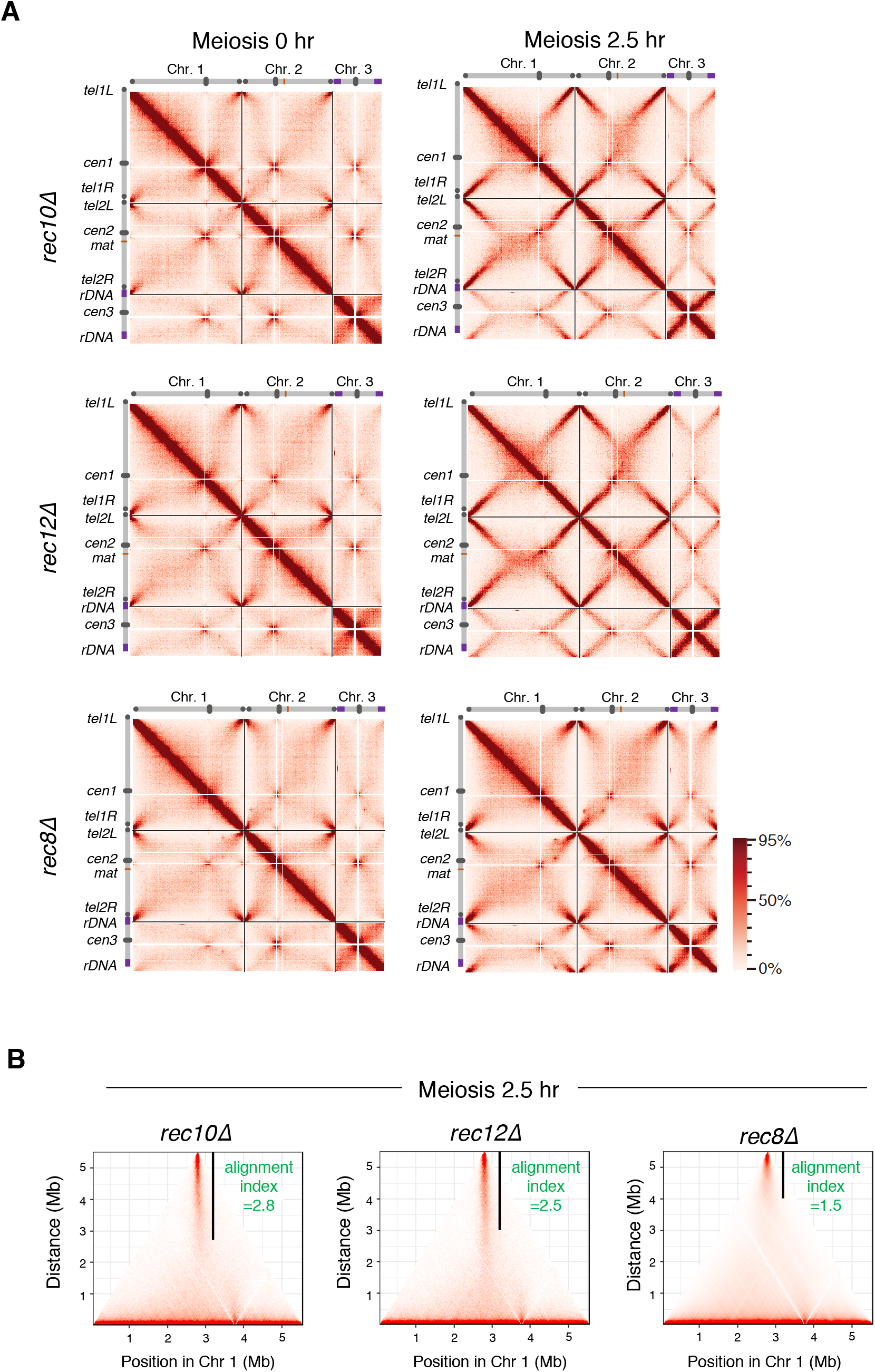
Rec8 cohesin complex shapes chromosomes into a structure suitable for its alignment. (A) Hi-C contact maps of three chromosomes for the indicated strains, prepared from the cells 0 and 2.5 h after entry into meiosis. The color scale on the right represents the percent ranking of contact scores. (B) The alignment index for the indicated strains at meiotic prophase (2.5 h after meiosis induction) is shown as the value along with a black vertical line in the Hi-C contact map of chromosome 1.

In the absence of Rec8, the bulk of the chromosomes do not follow the horsetail movement at meiotic prophase while only telomere regions follow the leading edge of the moving nucleus (39,84). This chromosome behavior reflects that the traction force generated by the horsetail movement is not transmitted to the chromosomes as proposed previously. The Hi-C results and chromosome movements suggest that the Rec8 cohesin complex forms the axis-loop chromatin structure and has structural properties to counteract the traction force (Figure 3F). This Rec8-dependent chromatin structure promotes the alignment of homologs without requiring LinE.

**Figure 3.**
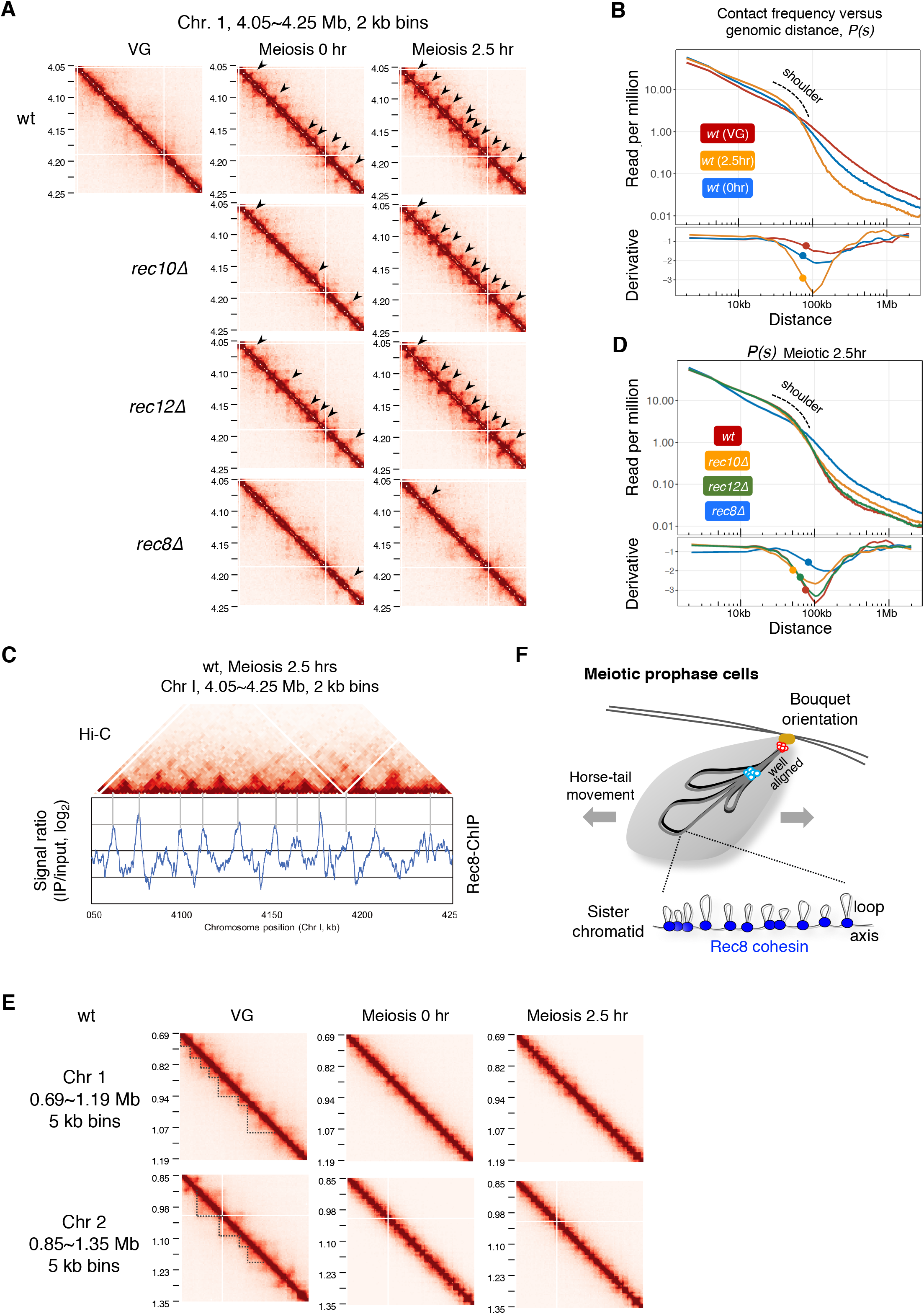
Rec8 cohesin complex is crucial to construct meiosis-specific chromatin loops. (A) Amplification of the Hi-C contact map in the 200 kb region of chromosome 1 plotted at 2 kb resolution for the indicated strains at 0 and 2.5 h after entry into meiosis (at VG only for wild-type). The black arrow depicts the loop structure. (B) Upper panel: Contact frequency versus genomic distance, *P(s)*, for VG (red) and meiosis 0 h (blue) and 2.5 h (orange) in the wild-type. The “shoulder” marked by the dotted line represents an increase in interactions within a region shorter than that length. Lower panel: The 1^st^-derivative of log-transformed Loess-smoothened curve is plotted. The filled circle indicates the transition point of the slope of *P(s)* obtained by 2^nd^-derivative. (C) A merged view of the Hi-C contact map and ChIP-chip data of Rec8 at the 200 kb region of chromosome 1 shown in (A). ChIP-chip data were adopted from a previously described source (85). (D) Upper panel: *P(s)*, for the wild-type (red), *rec8Δ* (blue), *rec10Δ* (orange), and *rec12Δ* (green) 2.5 h after meiotic induction. Lower panel: The 1^st^-derivative of log-transformed Loess-smoothened curve is plotted. The filled circle indicates the transition point of the slope of *P(s)* obtained by 2^nd^-derivative. (E) Amplification of Hi-C contact maps at the 500 kb region of chromosome 1 and 2 plotted at 5 kb resolution for the wild-type strain at VG, meiosis 0 h and 2.5 h. The dotted line depicts the TAD-like structure. (F) Schematic diagram of chromosome conformation at meiotic prophase in *S. pombe*. All chromosomes are tightly aligned from the telomeres to the centromeres. Rec8 establishes the meiosis-specific chromatin loop structure, irrespective of LinEs or recombination.

### Rec8 cohesin complex is crucial to form meiosis-specific chromatin loops

Magnification of Hi-C data (2 kb binning) in chromosome arm regions revealed that punctate Hi-C interactions occurred at the onset of meiosis (0 h) and became more prominent as meiosis progressed (2.5 h) in the wild-type (arrowheads in Figures 3A and S3A). As meiosis progressed in the wild-type, *P(s)* (the contact frequency versus distance) indicated a “shoulder,” representing an increase in interactions within a region shorter than that length (Figure 3C). The positions at the bases of these interactions coincided with previously identified Rec8 accumulation sites (85) (Figures 3B and S3B), suggesting that these punctate Hi-C interactions are generated by Rec8. In fact, punctate Hi-C interactions observed in the wild-type were mostly lost in *rec8Δ* (Figure 3A). In contrast, punctate Hi-C interactions were retained in *rec10Δ* or *rec12Δ* (Figures 3A and S3A). Consistently, the shoulder in *P(s)* was lost in the *rec8Δ* mutant but not in *rec10Δ* or *rec12Δ* (upper panel in Figure 3D). This difference in these mutants is also seen in the *P(s)* derivative plot (lower panel in Figure 3D). In the Hi-C data with 5 kb binning, topologically associating domain (TAD)-like signals were detected in wild-type VG cells (dotted lines in Figure 3E), representing the manifestation of relatively long-range interactions in mitotic chromosomes (27,64,80,86). As these TAD-like signals disappeared, Rec8-dependent loop structures appeared during meiosis (Figure 3E), similar to mammalian spermatocytes (80,87,88). These results suggest that a Rec8-dependent *cis*-loop structure is formed upon entry into meiosis in the wild-type, leading to the linear compaction of chromosomes (Figure 3F).

### Wpl1 in *S. pombe* regulates the length of chromatin loops in meiotic prophase

In somatic cells, the Rad21/Scc1 cohesin complex organizes chromatin with TADs, and Wapl depletion lengthens the chromatin loops and thickens the chromatin axis (34,37). However, evidence for Wapl affecting Rec8-dependent chromatin loops during meiosis remains elusive. To reveal involvement of Wapl in meiosis, we analyzed the chromosome structure in *wpl1Δ* cells. Fluorescence images of Rec8-GFP showed that axial structures in meiotic chromosomes were more prominent in *wpl1Δ* than in the wild-type (Figure 4A), suggesting that the axial structure becomes thicker in *S. pombe* during meiosis. To further examine the chromosome structures in *wpl1Δ*, we performed Hi-C analysis. We first confirmed synchronous progression of meiosis in *wpl1Δ* as in wild-type (Figure S4A). Hi-C contact maps showed that the background in *wpl1Δ* (upper panel of Figure 4B) was reduced than that in the wild-type (lower panel of Figure 4B). This reduced background probably reflects a reduction of the inter-and intra-chromosomal long-range interactions, including X-shaped contacts, in *wpl1Δ*. This notion is supported by fluorescence microscopic observations showing that alignment of chromosome arms was defective in *wpl1Δ* (Figure S4B) and that the chromosomes in *wpl1Δ* rarely showed torsional turning (Figure S4C, Supplemental Movies 1, and 2) during horsetail movements that is important for the alignment of homologs (89). These results suggest that Wpl1 plays a role in alignment of homologs through Rec8-dependent formation of axis-loop chromatin structure.

**Figure 4.**
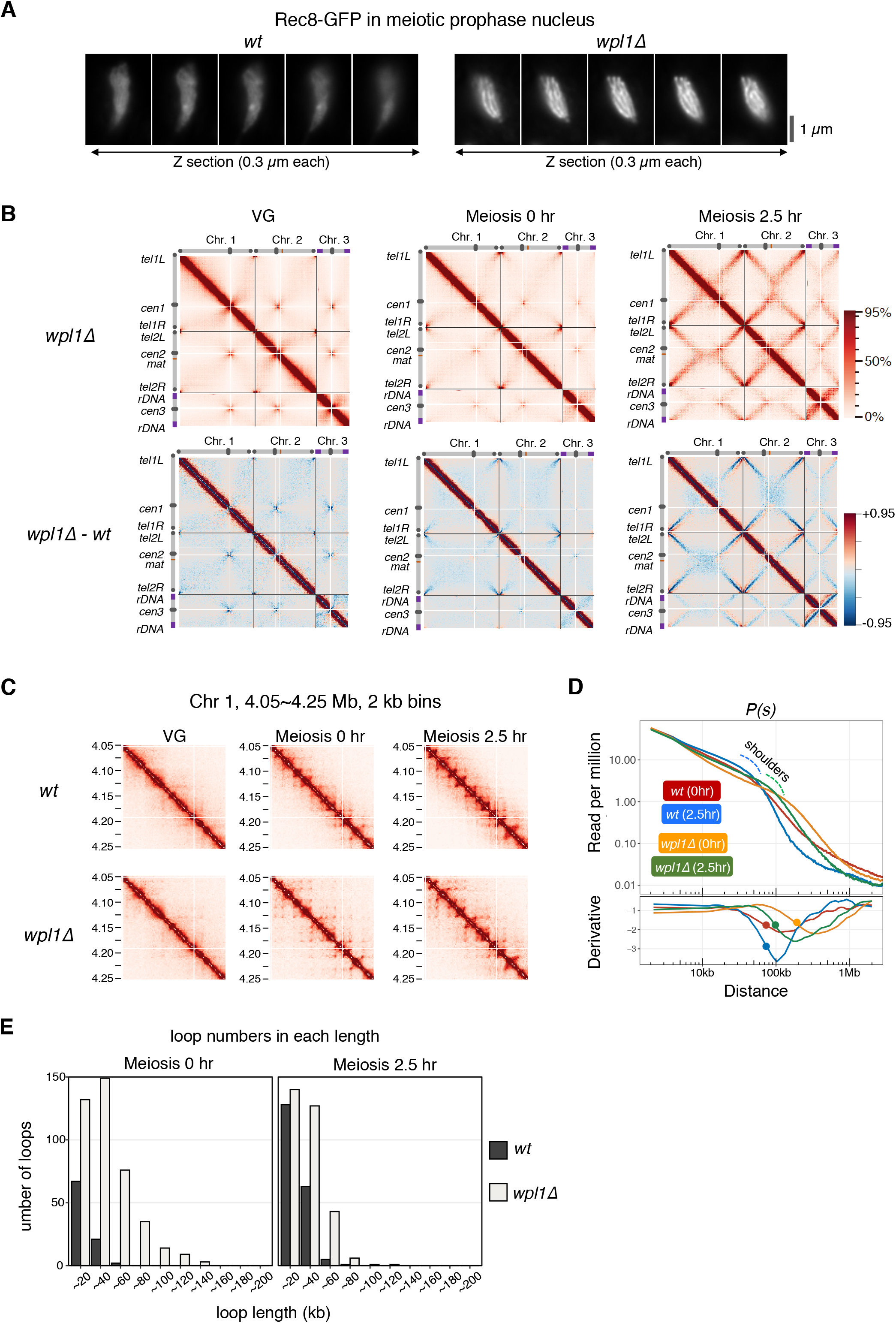
Wpl1 regulates chromatin loop length during meiotic prophase in *S. pombe*. (A) Serial Z sections (0.3 µm each) of Rec8-GFP signals at meiotic prophase in wild-type and *wpl1Δ*. Axis-like signals of Rec8 are clearly visible in *wpl1Δ*. (B) Upper panel displays Hi-C contact maps of three chromosomes at 20 kb resolution in *wpl1Δ* at vegetative growth (VG) and 0 and 2.5 h after entry into meiosis, with schematic views of each chromosome. The color scale on the right represents the percent ranking of contact scores. The lower panel displays heat maps showing the difference of contacts detected in each genome region by Hi-C in wild-type and *wpl1Δ* at VG and 0 and 2.5 h after entry into meiosis. The color scale on the right represents the percent ranking of difference scores. (C) Amplification of Hi-C contact maps showing the 200 kb region of chromosome 2 plotted at 2 kb resolution for the indicated strains at VG and 0 and 2.5 h after entry into meiosis. (D) Upper panel: *P(s)*s for meiosis 0 h and 2.5 h of wild-type (red and blue) and *wpl1Δ* (orange and green). The blue or green dotted line represents a shoulder of wild-type or *wpl1Δ* at 2.5 h after meiosis entry, respectively. Lower panel: The 1^st^-derivative of log-transformed Loess-smoothened curve is plotted. The filled circle indicates the transition point of the slope of *P(s)* obtained by 2^nd^-derivative. (E) Number of loops per length quantified using genome-wide loop caller HiCCUPS for wild-type (red, *wt*) and *wpl1Δ* cells (blue) at meiosis 0 h (left) and 2.5 h (right).

In the amplified view of Hi-C data (2 kb binning) in *wpl1Δ*, the number of grid-like punctate signals increased and the size of the diagonals of each grid increased upon entry into meiosis (Figures 4C and S4D), suggesting that loop lengths are longer in *wpl1Δ*. In fact, compared to the wild-type, the shoulder in *P(s)* showed a rightward shift in *wpl1Δ* (upper panel in Figure 4D). The derivative plot of *P(s)* also showed a rightward shift in *wpl1Δ* (lower panel in Figure 4D). Consistently, the HiCCUPS analysis showed that the number of longer loops increased in *wpl1Δ* (Figure 4E). These results imply that the proximal chromatin loops fuse with each other at their bases to form longer loops in the absence of Wpl1, leading to the linear compaction of chromosomes with thicker axes (Figure S4E).

Fluorescence microscopy showed that the thick Rec8 axis observed in *wpl1Δ* was maintained in *wpl1Δ rec10Δ* cells, in which LinE formation was lost (Figure S4F). Moreover, the thick Rec8 axis in the *wpl1Δ* background was formed in the mutants of Hop1 (a homolog of HORMAD) and Mek1 (Figure S4G), which are required for LinE formation (90). The thick Rec8 axis in the *wpl1Δ* background was also observed in the mutants of Rec12, Rec7, Rec15 (Figure S4G), and Mei4 (Figure S4H), which are required for recombination (91-94). These results further support our idea that the Rec8 axis formation is independent of LinE and recombination.

### Identification of *rec8* mutants showing defects in axis formation

Ectopic expression of Rad21 in meiosis partly compensates for the cohesion defect in the absence of Rec8 (57,59). However, Rad21 does not form a visible axis structure along meiotic chromatin, unlike Rec8 (Figure S5A). Therefore, besides the function of cohesion, Rec8 is expected to have an additional function that is more robust than that of Rad21 in chromatin axis formation. Based on this idea, we attempted to identify mutants of *rec8* that specifically shows defects in axis formation without affecting its cohesion function. To this end, we constructed the *S. pombe* strain in which Rec8 and Rec11 were ectopically expressed as the sole source of cohesin during mitosis, replacing Rad21 and Psc3 (57) (Figure 5A). Using this strain as a screening host, randomly mutagenized *rec8* cDNAs fused with GFP-coding DNA were transformed into the host strain. Since Rec8 is the only available kleisin protein in this strain, transformants bearing a *rec8* mutation with a loss of cohesion show a lethal or slow-growth phenotype. Furthermore, we used the *h*^*90*^ mating-type in the *mei4Δ* background as a host strain to arrest each transformant at the meiotic prophase on agar plates with the sporulation medium (91). We also introduced *wpl1Δ* in the host strain to produce a prominent axis of Rec8-GFP (Figure 4A) so that defects in axis formation could be easily evaluated by microscopic observation (Figure 5A).

**Figure 5.**
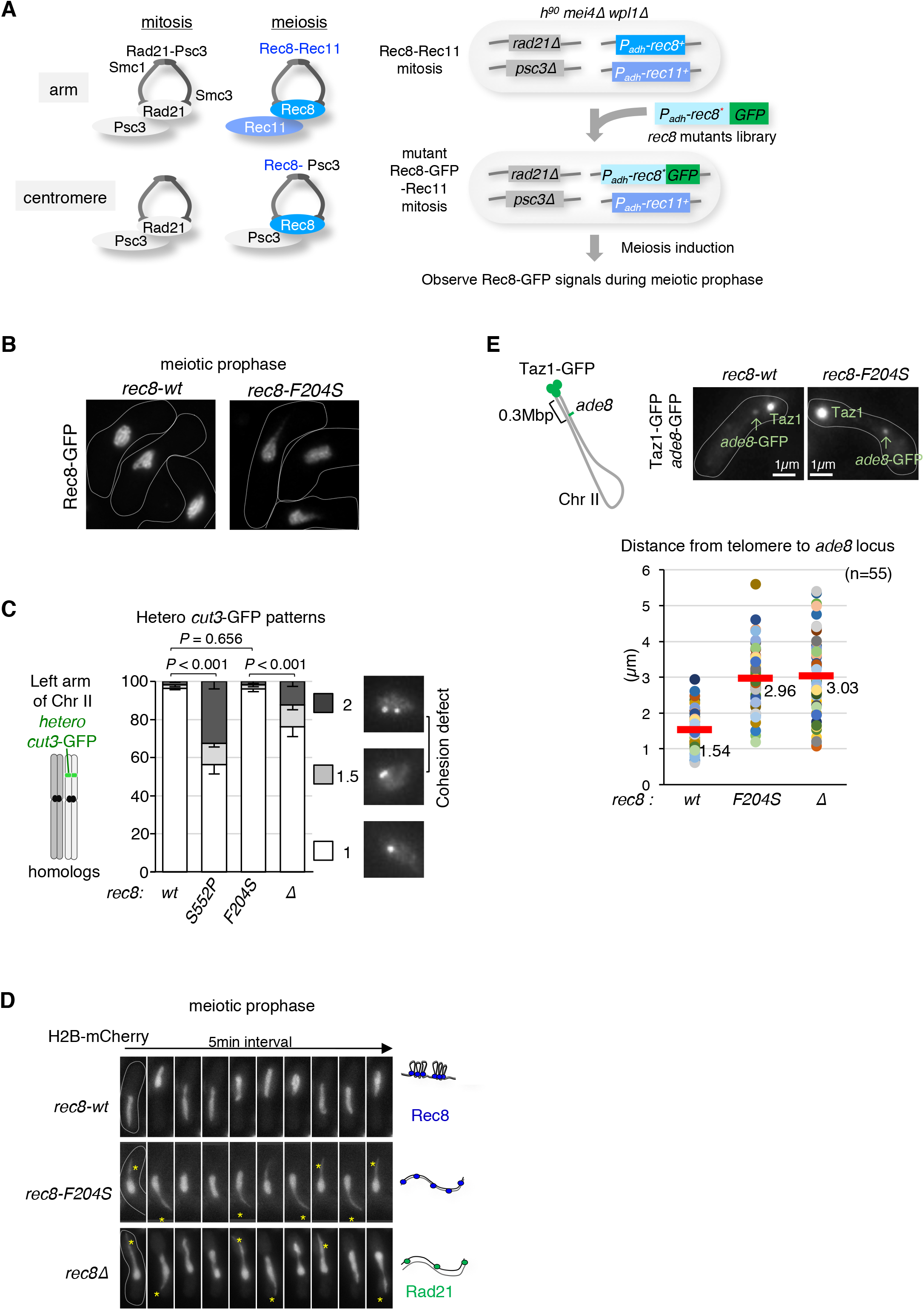
Isolation of a *rec8* mutant showing a defect in axis formation with an intact cohesion function. (A) The left panel shows a schematic view of cohesin complexes in fission yeast in mitosis and meiosis. In mitosis, the Rad21-Psc3 cohesin complex is ubiquitously located along the entire chromosome. In meiosis, the Rec8-Psc3 cohesin complex exists only at the centromeres, while the Rec8-Rec11 cohesin complex is present at the chromosomal arm regions. The right panel shows the scheme for isolation of *rec8* mutants that specifically show defects in axis formation but display normal cohesion in meiotic prophase. The promoter of the *rec8* gene is replaced by the *adh1* promoter, and the promoter of the *rec11* gene is replaced by the *adh41* (a weaker version of *adh1*) promoter, to allow gene expression both in mitosis and meiosis. After transformation of a library of GFP-fused *rec8* mutants with a selective marker, *bsdR* (not shown here), the blasticidin-resistant colonies obtained were subsequently induced to undergo meiosis to observe the axial Rec8-GFP signals. Growth rate of the blasticidin-resistant clone acts as an indicator of the functionality of Rec8-Rec11 cohesin. (B) Representative images of Rec8-GFP depicting cell shape (white dotted line) during meiotic prophase in wild-type and the isolated *rec8-F204S* mutant in this screening. (C) Cohesion defects in the chromosome arm region (*cut3*^*+*^ locus, left arm of chromosome 2) were examined in the indicated cells arrested at prophase I by the *mei4Δ* mutation. After fixation by methanol, the number of heterozygous *cut3*-GFP signals was counted (n > 150). A score of 2 or 1.5 *cut3*-GFP signals represents a complete or partial loss of sister chromatid cohesion, respectively. The *rec8-S552P* identified in this screening displays a remarkable defect in cohesion that is stronger than that of *rec8Δ* (see the text). Error bars show standard deviations from three independent experiment. P values were obtained from Chi-square test for statistical significance. (D) Time-lapse observations (5 min intervals) of horsetail nuclear movement in the indicated cells stained with histone H2B-mCherry. The cartoon on the right represents a model of the chromatin state for each strain. In *rec8Δ*, Rad21 partially compensates for the cohesion defect. (E) The upper left panel shows a schematic diagram of the distance of *ade8*-*lacO* inserts from the telomere of chromosome II. The upper right panels represent images of GFP signals for the indicated strains. Bright GFP signals at the edge of the nucleus represent telomeres (Taz1-GFP), and weak GFP signals represent the *ade8* locus. The graph at the bottom displays a plot of the distance between the telomere and the *ade8* locus in each cell during meiotic prophase for the indicated strains. The average distance is plotted (n = 55).

Among approximately 3,000 viable transformants of the *rec8* mutant library, we searched for mutants that showed axis formation defects with normal cohesion function under a fluorescence microscope. We identified a mutant that displayed normal growth (Figure S5B) but lost the Rec8 axis structure (Figure 5B). In this mutant, the amino acid phenylalanine at the 204^th^ residue of Rec8 was substituted with serine (*rec8-F204S*). The *rec8-F204S* mutant maintained sister chromatid cohesion as assessed at the *cut3* gene locus (Figure 5C) and other chromosomal loci (Figure S5C); *rec8-S552P* and *rec8Δ*, which showed the cohesion defect, were used as a control strain (see Figure S5C, S5D, and S5E for details of the *rec8-S552P* mutant). Moreover, a normal cohesin complex was formed in the *rec8-F204S* mutant, with comparable expression of Rec11 (Figures S5F and S5G).

To delineate the axis formation defects caused by *rec8-F204S*, we observed the chromosome morphology during horsetail movement in meiotic prophase. In the *rec8-F204S* mutant, only the leading edge of the nucleus followed the horsetail movement, while the bulk of chromosomes were left behind, similar to *rec8Δ* (Figure 5D, Supplemental Movies 1, and 3). To determine the degree of chromosome compaction more directly, we measured the distance between the telomere and the *ade8* gene locus during meiotic prophase using a GFP-tagged telomere protein (Taz1) and a *lacO*/lacI-GFP tag at the *ade8* gene locus (Figure 5E). As in *rec8*Δ, the telomere-*ade8* distance was longer in the *rec8-F204S* mutant than in the wild-type, suggesting that the chromatin of the *rec8-F204S* mutant was flexible and was abnormally stretched by the traction of the horsetail movement.

To evaluate the specificity of the amino acid at the 204^th^ residue of Rec8 in terms of the axis formation, we examined Rec8 mutants bearing other amino acid substitutions. F204T, F204N, F204D, F204A, F204R, and F204H mutants showed defects in the Rec8 axis formation, whereas F204C, F204W, and F204L mutants formed the normal Rec8 axis (Figure S5H). These results suggest that the hydrophobicity of this residue in Rec8 may be important for its axis-forming function.

### *rec8-F204S* mutant is defective in chromatin loop formation

To examine the chromatin loop formation in the *rec8-F204S* mutant defective in axis formation, we performed Hi-C analysis in the *rec8-F204S* mutant as performed for the *rec8-wt* (Figure S6A). In the Hi-C contact maps of three chromosomes at meiotic prophase (2.5 h), formation of X-shaped patterns representing the bouquet orientation were evidently impaired in the *rec8-F204S* mutant, as in *rec8Δ* (Figure 6A). The alignment index of chromosome 1 at meiotic prophase (2.5 h) decreased in *rec8-F204S* (1.6) compared with the wild-type (*rec8-wt*, 2.6) (Figure 6B). Moreover, in the Hi-C data with amplified 2 kb binning, the number of punctate Hi-C interactions decreased in *rec8-F204S* (Figure 6C), although it was not as prominent as in *rec8Δ*. In addition, the shoulder in *P(s)* became flattened in *rec8-F204S* compared to *rec8-wt*, both at 0 and 2.5 h after meiosis induction (Figure 6D). Consistently, chromatin loops with a length of approximately 20 kb that formed in a Rec8-dependent manner were diminished at the meiotic prophase in *rec8-F204S* (Figure 6E). These results suggest that in the *rec8-F204S* mutant, as in *rec8Δ*, the Rec8-dependent meiosis-specific short chromatin loop structures are lost, resulting in a concomitant loss of the structural property of the chromosome required for proper alignment.

**Figure 6.**
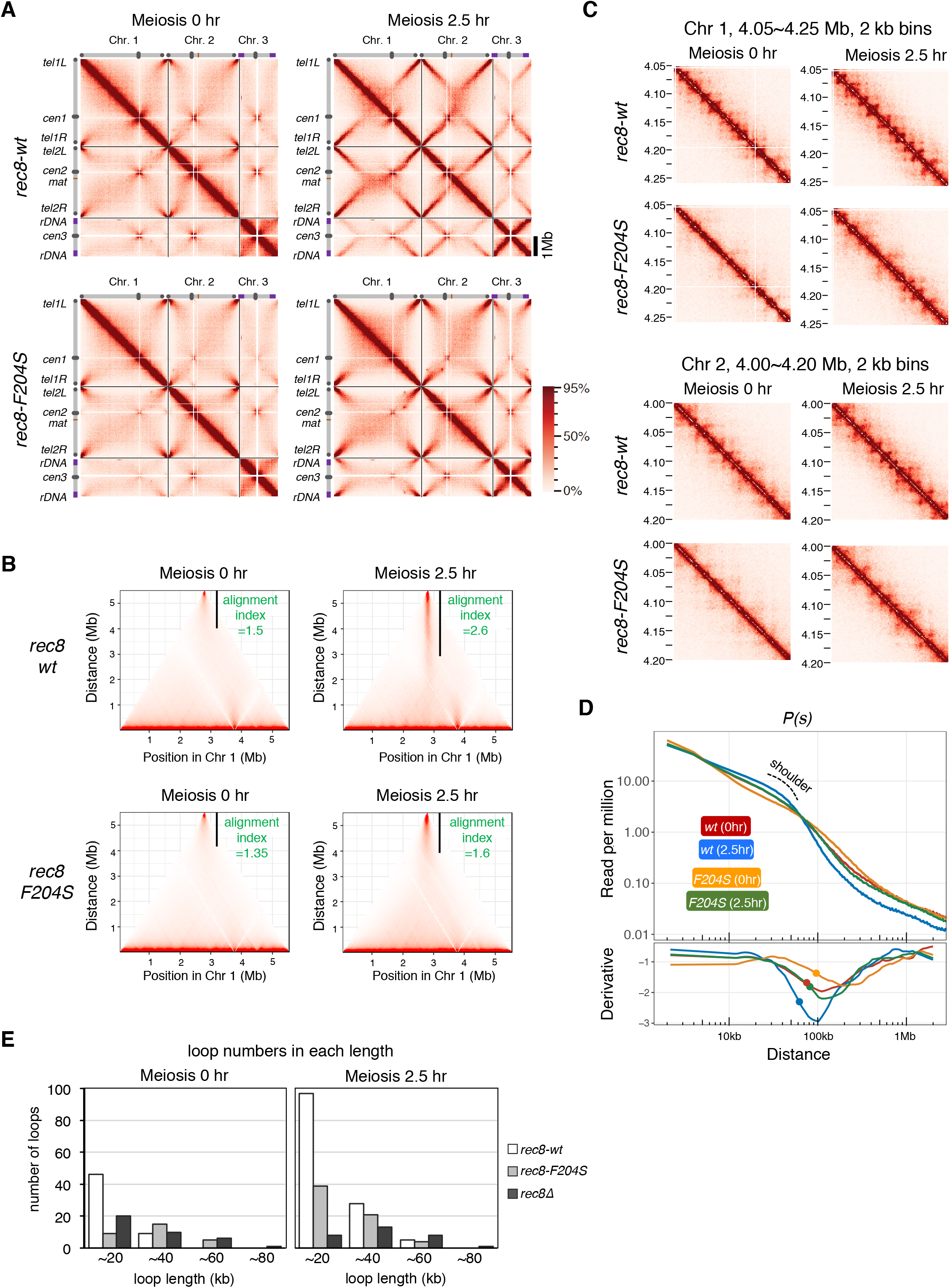
The *rec8-F204S* mutant displays defective chromatin loop formation. (A) Hi-C contact maps of three chromosomes at 20 kb resolution in wild-type *rec8*-3×Pk and *rec8-F204S*-3×Pk cells at 0 and 2.5 h after entry into meiosis, with schematic views of each chromosome. The color scale on the right represents the percent ranking of contact scores. (B) The alignment index for indicated strains at 0 and 2.5 h after meiosis induction is shown as the value along with a black vertical line in the Hi-C contact map of chromosome 1. (C) Amplification of Hi-C contact maps of the 200 kb region of chromosome 1 (upper panel) and chromosome 2 (lower panel) at 2 kb resolution, for the indicated strain at 0 and 2.5 h after entry into meiosis. (D) Upper panel: *P(s)*, for 0 and 2.5 h after entry into meiosis in *rec8-wt* (red and blue) and *rec8-F204S* (orange and green). A shoulder observed at 2.5 h after entry into meiosis for *rec8-wt* is shown with a dotted line. Lower panel: The 1^st^-derivative of log-transformed Loess-smoothened curve is plotted. The filled circle indicates the transition point of the slope of *P(s)* obtained by 2^nd^-derivative. (E) Number of loops per unit length quantified using genome-wide loop caller HiCCUPS for the indicated strains during meiosis at 0 h (left) and 2.5 h (right).

Next, we performed a chromatin immunoprecipitation (ChIP) assay to determine the localization pattern of Rec8 on chromatin. The accumulation of Rec8-F204S protein apparently decreased compared with wild-type Rec8 at the loci of known cohesin-enriched regions (Figure S6B). In contrast, the localization in other chromosome arm regions seemed to be normal (Figure S6B). Rec8-enriched regions mostly coincided with each base of the chromatin loops (Figures 3C and S3B). In the case of *rec8-F204S*, the amount of Rec8 in the non-enriched region appeared normal. Thus, the reduced accumulation of Rec8 at known enriched regions possibly reflects a noticeable loss of chromatin loop formation (Figure S6C).

### *rec8-F204S* mutant is defective in LinE formation and recombination

Next, we examined the formation of LinEs in the *rec8-F204S* mutant. In wild-type cells, the LinE component Rec10-mCherry localized within the entire nucleus, similar to Rec8-GFP (Figures 7A and S7A; *CK1*^*+*^, *rec8*^*+*^, *rec11-wt*). In *rec8-F204S* cells, however, Rec10-mCherry formed aberrant dotty or filamentous aggregates within the nucleus, similar to *rec8Δ* and *rec11-5A* cells in which LinE was completely lost (Figures 7A and S7A) (48). Consistent with the LinE formation defect, *rec8-F204S* cells showed a decrease in the frequency of crossover recombination at several loci (Figure 7B). Intragenic recombination between the genomic *ade6-M26* allele and the plasmid harboring the *ade6-469* allele was also significantly reduced in *rec8-F204S* (Figure S7B), similar to *rec8Δ*. Rec11 phosphorylation by CK1, which is required for the assembly of LinEs (48), occurred normally in the *rec8-F204S* mutant, similar to the wild-type (Figure 7C). Expression of a phospho-mimetic mutant of Rec11 (*rec11-5D*), which can suppress the defects caused by the loss of Rec11 phosphorylation via CK1 (48), did not suppress defects in LinE formation or meiotic recombination in *rec8-F204S* (Figures S7A and S7C). These results suggest that axis-loop formation along sister chromatids via the Rec8 cohesin complex is still essential for LinE assembly, even in the presence of phosphorylated Rec11. Collectively, these results demonstrate that Rec8-dependent axis-loop structures facilitate meiotic recombination through the LinE assembly (Figures 7D).

**Figure 7.**
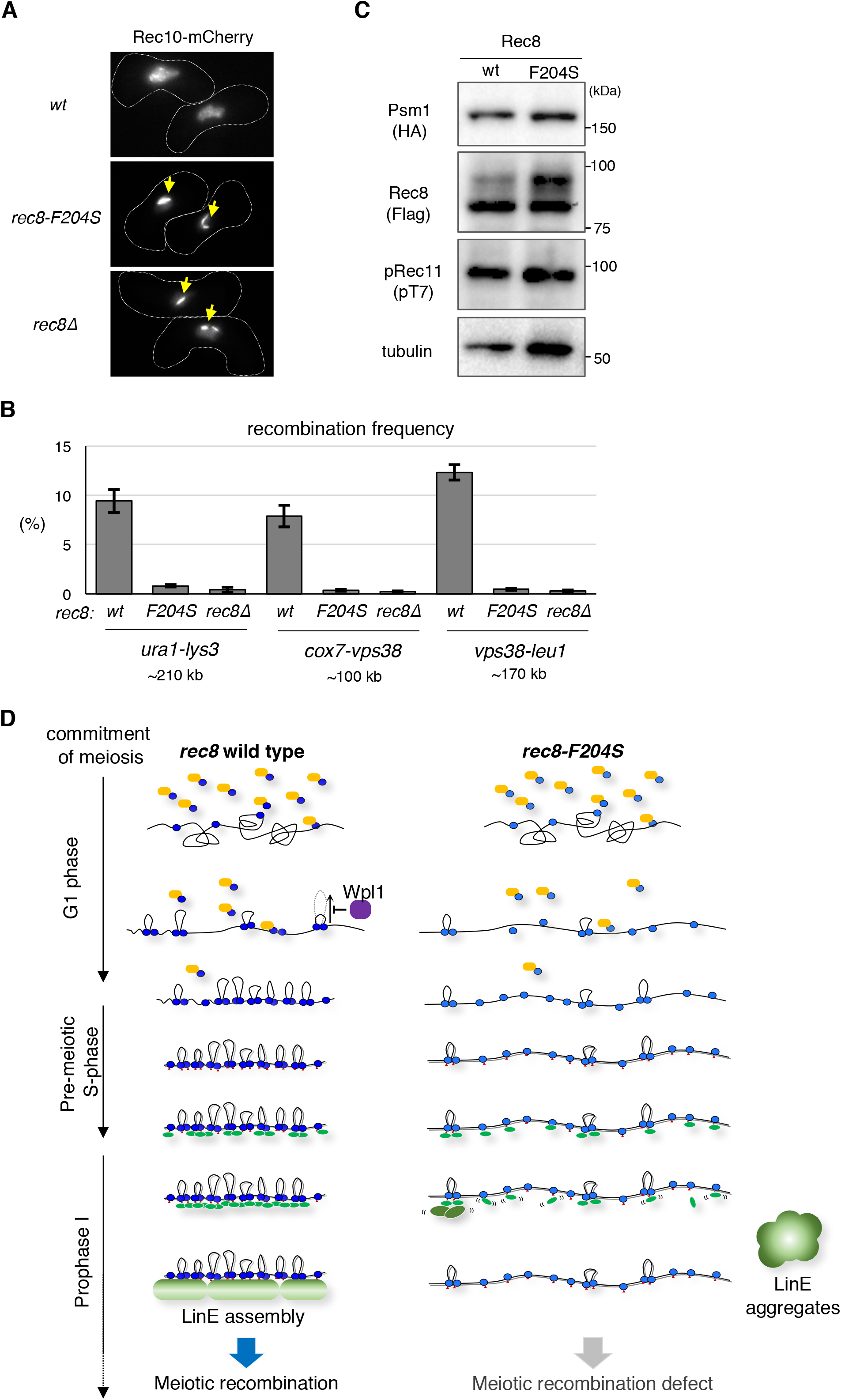
Rec8-dependent chromatin loop formation and meiotic recombination. (A) Representative pictures of Rec10-mCherry depicting cell shape (white dotted line) during meiotic prophase for the indicated strains. The yellow arrows depict the aggregates of LinE. (B) Intergenic meiotic recombination for *ura1-lys3* (n > 700), *cox7-vps38* (n > 500), and *vps38-leu1* (n > 500) in the indicated zygotes. Error bars show standard deviations from three independent experiments. (C) Western blot analysis. Whole-cell extracts were prepared from cells in meiotic prophase (2.5 h after meiosis induction) expressing Rec8-3×Flag in wild-type and *rec8-F204S*, with Psm1-3×HA. Rec11 phosphorylation by CK1 was detected using anti-phosphorylated Rec11 (at T7). The monoclonal anti-tubulin (DM1A, Abcam) antibody was used as a loading control. (D) A model for the Rec8-dependent regulation of LinE assembly and meiotic recombination. After the commitment to meiosis, the Rec8 cohesin complex (shown in blue) is loaded onto the chromosomes via the cohesin loader complex (Mis4-Ssl3) (shown in orange). Once loaded, the Rec8 cohesin complex promotes loop formation, which is antagonized by Wpl1 (shown in purple). By meiotic prophase, in parallel with the continuous loading of Rec8 and phosphorylation of Rec11 by CK1 (indicated by red triangles), chromatin looping with axis formation by the Rec8 cohesin complex occurs to form chromatin to recruit LinE factors. Refer to the detailed description in the Discussion.

## DISCUSSION

In this study, we have demonstrated the Rec8-dependent formation of axis-loop structures during meiosis in *S. pombe*, an asynaptic organism. The Rec8-dependent axis-loop structure functions as a scaffold for the formation of the LinE axis, thereby ensuring homologous recombination during meiosis.

### Pairing of homologous chromosomes in *S. pombe* meiosis

Our Hi-C analysis revealed the emergence of an intriguing X-shaped contact map representing tight contacts between chromosome arm regions near the telomere in the meiotic prophase of *S. pombe*. This result is consistent with previous microscopic observations of telomere clusters at the SPB (the bouquet orientation of chromosomes) during meiotic prophase (75,79) (Figure 3F). The Hi-C data indicate that contacts occur not only between homologous arms but also between non-homologous arms. It should be noted that chromosomes recognize their homologous pair among a bundle of all the chromosomes. Therefore, mechanisms to eliminate non-homologous pairing while simultaneously promoting homologous pairing may exist. Such selective forces between chromosomes can be provided by the oscillatory movement of telomere-bundled chromosomes (89,95). Additionally, noncoding RNAs play a robust role in recognizing homologous pairs (76,78,96). Importantly, the Rec8-dependent formation of axis-loop chromatin structure is necessary for optimizing these mechanisms involving telomere clustering, horsetail movement, and noncoding RNA-mediated recognition.

### Possible mechanisms for formation of the Rec8 axis-loop chromatin

A loop extrusion model has been proposed to form chromatin loops by SMC protein complexes including cohesin (26,28,30,97). In this model, the SMC protein complex interacts with DNA, reels, and extrudes it through its ring structure to form a DNA loop (98-102). In addition, *cis*-looping of distal chromatin or TAD formation is thought to be established by the action of the Rad21 cohesin complex anchored on chromatin via the CCCTC-binding factor (CTCF) in mammalian somatic cells (66,103-105). CTCF is conserved in vertebrates (106). In *S. pombe*, CTCF is not present, and proteins that replace the function of CTCF are unknown. In contrast, Scc2 and Scc4 are conserved from yeasts to humans (Mis4 and Ssl3 in *S. pombe*, respectively); human Scc2 and Scc4 are required for the DNA extruding activity of human Rad21 cohesin complex at least *in vitro* (98). In *S. pombe*, chromatin loop structures are already formed at the onset of meiosis (0 h) when the Rec8 cohesin complex is loaded on the chromatin by Mis4 and Ssl3 before pre-meiotic DNA replication (Figures 3A and S1A). Thus, Mis4 and Ssl3 can be initial factors that create chromatin loops in *S. pombe* meiosis. Alternatively, some changes in chromatin potential, involving histones, Rec8, or other yet-unidentified factors, may occur upon entry into meiosis to form the chromatin axis-loop structure.

The *rec8-F204S* mutant isolated in this study showed a significant defect in the formation of axis-loop chromatin structures, despite maintaining the cohesion function. The F204 residue of Rec8 locates in the kleisin-Scc2 interaction domain, as previously shown by an *in vitro* binding assay (107). However, this phenylalanine residue is not conserved in vertebrates (Figure S5I) despite conservation in neighboring sequences. Thus, Rec8 and Scc2/Mis4 with Scc4/Ssl3 may also play some roles in modulating meiosis-specific chromatin loop structures, although precise mechanisms remain to be elucidated.

### Rec8 axis-loop structure as a scaffold for LinE assembly

In the *rec8-F204S* mutant, which was unable to form the axis-loop structure, LinE formation failed to occur, and the recombination frequency was markedly reduced (Figures 7A, S7A, 7B, and S7B). However, phosphorylation of Rec11, which is required for LinE formation, was normal in the *rec8-F204S* mutant (Figures 7C), suggesting that mechanisms other than Rec11 phosphorylation are involved in the LinE defects of this mutant. It has been reported that chromosomal localization of Rec8 does not exactly overlap with that of Rec10 (a LinE component) despite the interaction of the Rec8 cohesin complex with Rec10 (48,92,108). These results imply that LinE factors, including Rec10, localize to chromosomes initially through interaction with the Rec8 cohesin complex and then redistribute along the chromosome appropriately to assemble LinEs. Based on these notions, we speculate that oligomerization of LinE factors requires the Rec8 axis as a scaffold, which is lost in the *rec8-F204S* mutant. Without this Rec8 scaffold, the association between LinE factors and chromatin would not be stabilized, resulting in the formation of aggregates (Figures 7A, S7A, and 7D), probably due to their propensity to adhere to each other. These aggregates are reminiscent of the polycomplex formed by mutants with defects in SC formation during meiosis of *S. cerevisiae* (2).

Expression of the Rec11-Rec10 fusion protein as well as the Rec11-5D mutant protein, both of which can suppress the defects caused by the loss of Rec11 phosphorylation (48), did not suppress the LinE formation defect of *rec8-F204S* (Figures S7A, S7C and S7D). These results support the notion that the defects in LinE formation in this mutant are due to the unstable interaction between Rec10 and the Rec8 cohesin complex (Figures 7D). LinEs in *S. pombe* are crucial in chromatin recruitment of factors that induce DNA double-strand breaks for recombination (92). This fact explains why the *rec8-F204S* mutant fails to undergo meiotic recombination. Although the F204 residue in mouse is leucine, the phenylalanine-to-leucine substitution in *S. pombe* Rec8 showed almost no defects in axis formation (Figures S5H and S5I). Rec8 and Rec11/SA3 are required for lateral element formation in mammals, as well as for LinE formation in *S. pombe* (18-21,46,48,49,83). Thus, there may be a mechanism by which meiosis-specific cohesin-dependent axial structures can function as a scaffold in the formation of lateral elements in mammals.

In conclusion, the current results obtained in *S. pombe* indicate that the Rec8-dependent axis-loop chromatin structures are crucial for meiotic recombination. As meiotic recombination is essential for faithful propagation of chromosomes into the progenies during meiosis in eukaryotes, this chromatin structure will provide a basis for understanding the molecular mechanisms of meiotic chromosome propagation in mammals, potentially related to miscarriages and congenital abnormalities in humans (4,5).

## DATA AVAILABILITY

Hi-C data were submitted to Gene Expression Omnibus (GEO) with the accession number: GSE192516. Further data are available from the corresponding authors upon reasonable request.

## SUPPLEMENTARY DATA

Supplementary Data are available at NAR online.

## ACKNOWLEDGEMENT

We thank Drs. Masamitsu Sato, Shigehiro Kawashima and Juraj Gregan for providing *S. pombe* strains, Dr. Kayoko Tanaka for the helpful advice to induce highly synchronous meiosis, and all members of our laboratory for their valuable support and discussion.

## FUNDING

This work was supported by JSPS KAKENHI grants JP26711020 (to T.S.), JP19K06503 (to D.Q.D.), JP18H05528 (to T.H.), JP20H00454 and JP18H05533 (to Y.H.), and also by the National Institutes of Health/National Institute of General Medical Sciences (R01GM124195 to K.N.).

## CONFLICT OF INTEREST

The authors declare no competing interests.

## Notes

### Competing Interest Statement

The authors have declared no competing interest.

### Summary of Updates

Main text and Supplemental files updated

