## Supplemental figures 1 to 7 and table 1 and 2 for "Rec8 cohesin-mediated axis-loop chromatin architecture is required for meiotic recombination"

### Supplementary Figure S1-S7

### Supplementary Table S1 and S2

### Legend of Supplementary Movie S1-S3

### Supplementary Figures

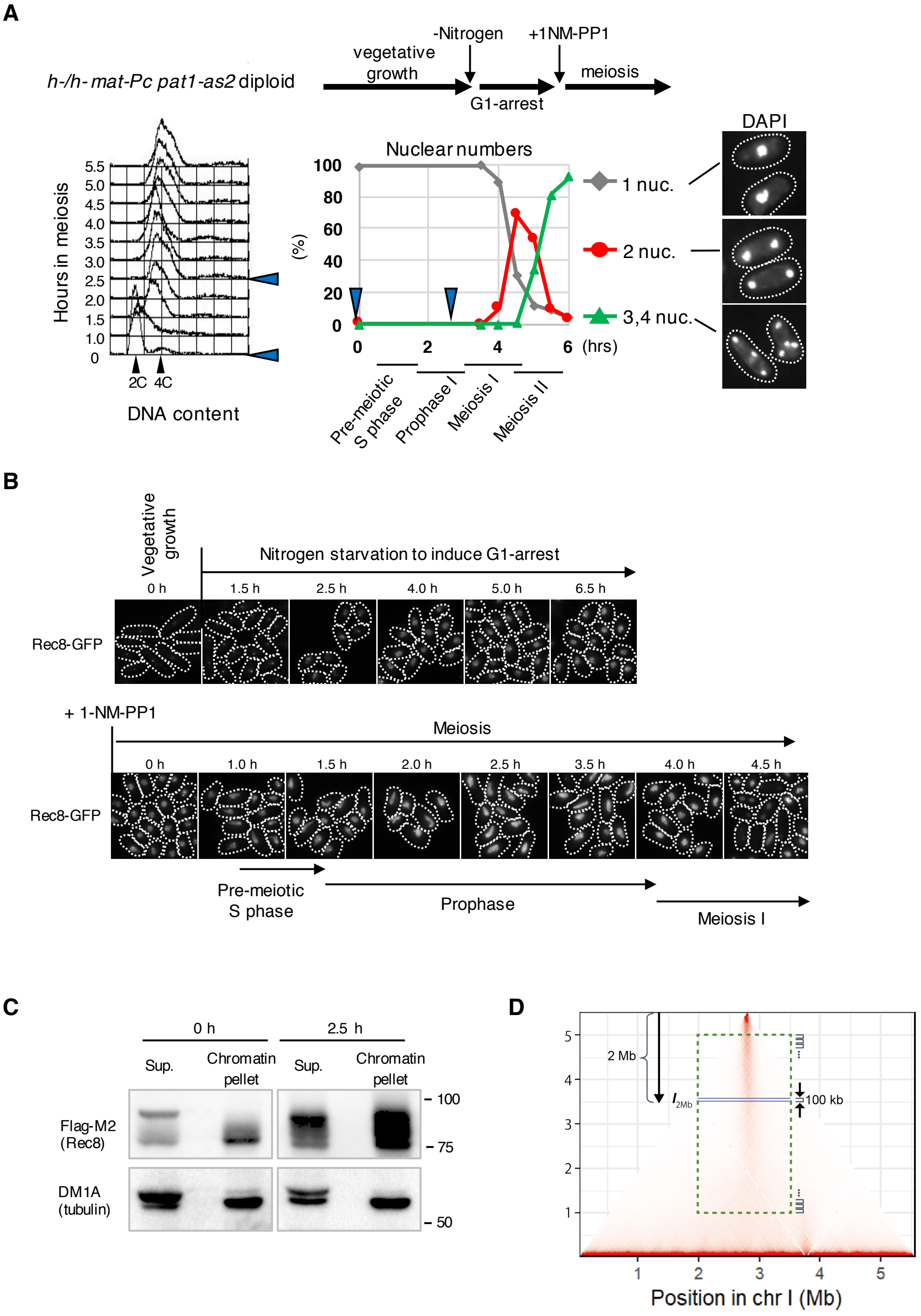


**Figure S1.** Preparation of synchronous populations of meiotic cells for Hi-C analysis (A) Top bar depicts the scheme for synchronous induction of meiosis using *h⁻/h⁻ pat1-as2 MDR-sup* cells. The left panel displays the meiotic progression of *h⁻/h⁻ pat1-as2 MDR-sup rec8⁺-GFP* wild type cells, monitored by fluorescence-activated cell sorting. The right panel displays the meiotic progression of *h⁻/h⁻ pat1-as2 MDR-sup rec8⁺-GFP* wild type cells, monitored by the quantification of the number of nuclei stained using

4′,6-diamidino-2-phenylindole (DAPI). Two nuclei indicate the cells in nuclear division of meiosis I; three of four nuclei indicate those in meiosis II. Representative pictures of the right-most cells. (B) Representative images of the Rec8-GFP with cell shape (white contour) at indicated time points. (C) Western blot of wild type cells expressing Rec8-3×Flag. At 0 h and 2.5 h after meiosis entry, whole-cell extracts were harvested in low salt buffer (150 mM NaCl). The cell extracts were centrifuged and the supernatants and pellets were subjected to western blotting using Flag-M2 monoclonal antibody (Sigma-Aldrich) to detect Rec8, and monoclonal anti-tubulin (DM1A, Abcam) as a loading control. (D) Calculation of the alignment index. ICE-normalized Hi-C contact scores at 20 kb resolution were scaled by the following calculation:

$$\boldsymbol{S_{i,j}} = \boldsymbol{N_{i,j}} \div \Sigma \boldsymbol{N_{i,j}} \times \text{(total combination number)}$$

where $\boldsymbol{S_{i,j}}$ is the scaled score of the $\boldsymbol{i}$ and $\boldsymbol{j}$ combination, and $\boldsymbol{N_{i,j}}$ is the ICE-normalized score of the $\boldsymbol{i}$ and $\boldsymbol{j}$ combination. The average scaled scores were set to 1. To calculate the alignment index, scaled scores were extracted from the area between 2–3.5 Mb and 1–5 Mb of chromosome 1 (green dotted square). The relative intensity ($\boldsymbol{I}$) was calculated for each 100-kb section ($\boldsymbol{w}$) as follows:

$$\boldsymbol{I_w} = \text{(average of top 5\% score)} - \boldsymbol{B},$$

where background $\boldsymbol{B}$ was calculated using the average of the bottom 70% scores. The alignment index was defined as the shortest distance from the telomere, where

$$\boldsymbol{I_w} - \text{(minimum of } \boldsymbol{I_w}) \geqq 0.02.$$

The alignment index corresponds to a length of the solid anti-diagonal line.

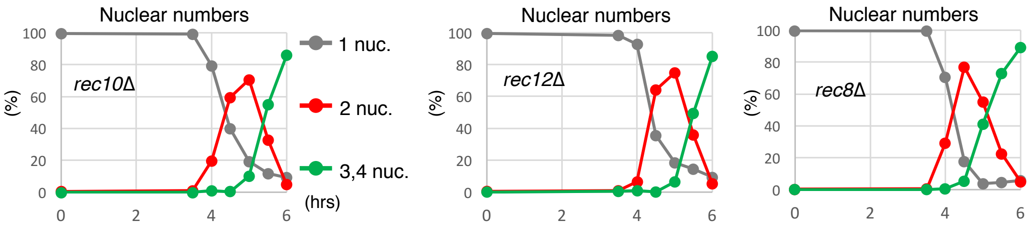


**Figure S2.** Synchronous progression of meiosis in the mutant strains
Synchronous progression of meiosis in the indicated mutant strains in the *h⁻/h⁻ pat1-as2 MDR-sup* background, monitored by the quantification of the number of nuclei stained with DAPI.

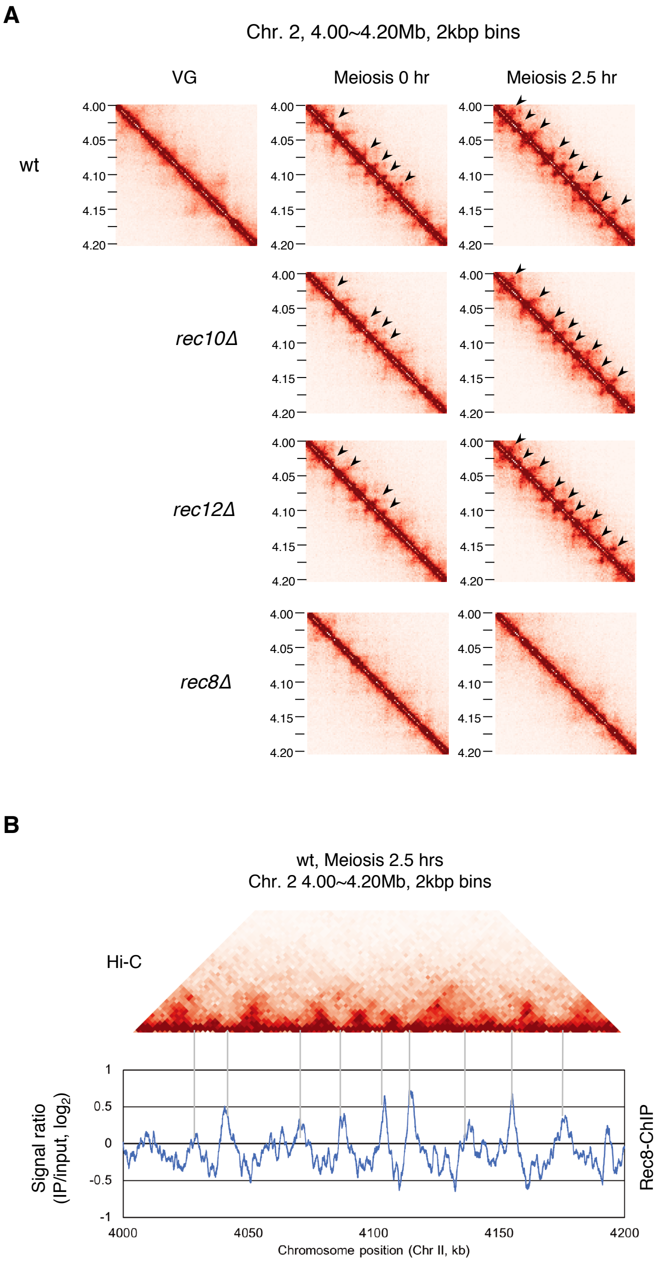


**Figure S3.** High resolution Hi-C analysis
(A) Amplification of Hi-C contact maps of the approximately 200 kbp region of chromosome 2 plotted at 2 kbp resolution for the indicated strains at meiosis 0 h and 2.5 h, and in VG only for wild type. The black arrows depict the loop structure. (B) A merged view of Hi-C contact map of a 200 kbp region of chromosome 1 used in (A) and ChIP-chip data of Rec8 at the same region (85).

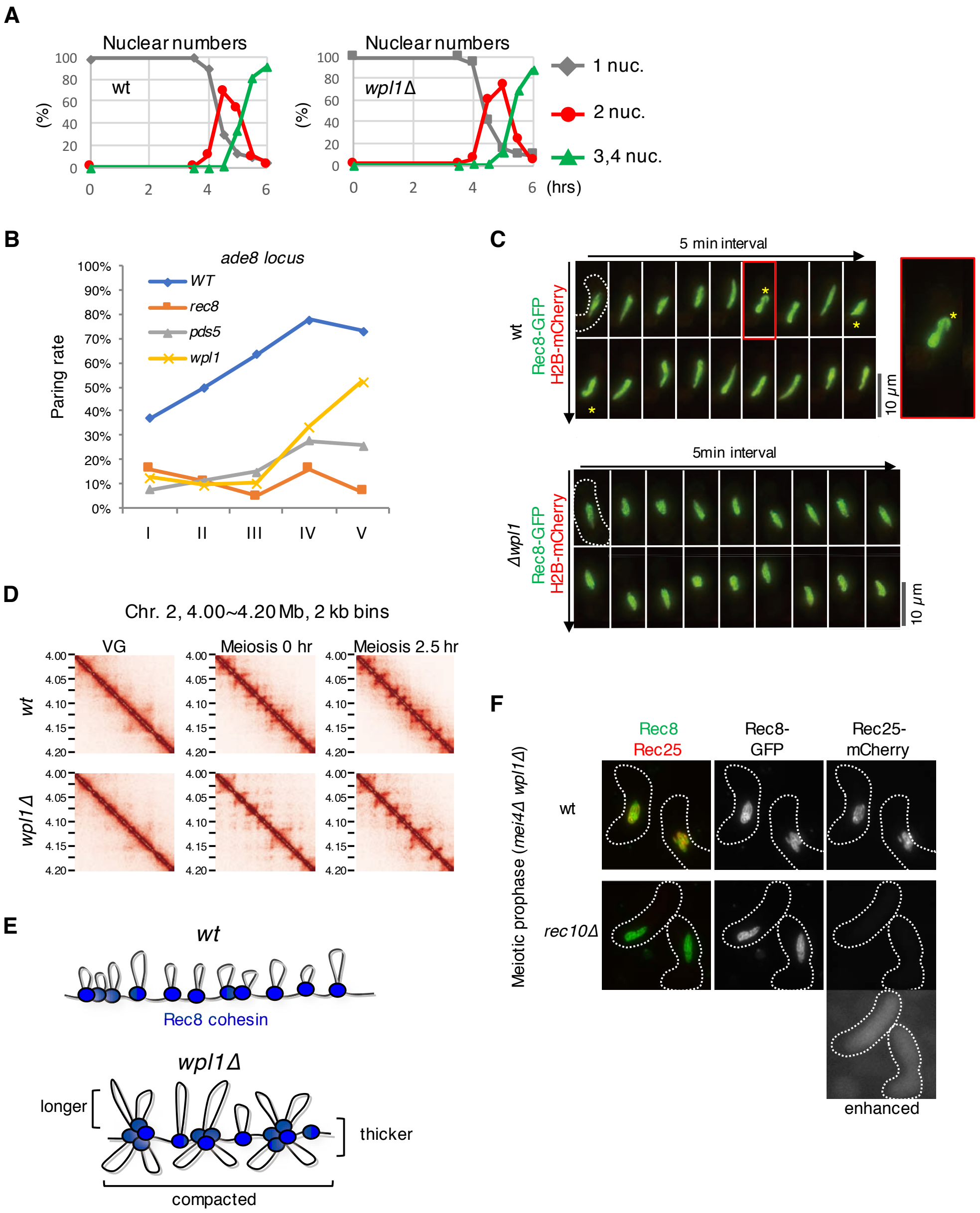
A
Nuclear numbers
wt
wpl1Δ
(%)
100
80
60
40
20
0
0
2
4
6
(hrs)
1 nuc.
2 nuc.
3,4 nuc.
B
ade8 locus
WT
rec8
pds5
wpl1
Paring rate
100%
90%
80%
70%
60%
50%
40%
30%
20%
10%
0%
I
II
III
IV
V
C
5 min interval
wt
Rec8-GFP
H2B-mCherry
10 μm
5min interval
Δwpl1
D
Chr. 2, 4.00~4.20 Mb, 2 kb bins
VG
Meiosis 0 hr
Meiosis 2.5 hr
4.00
4.05
4.10
4.15
4.20
E
Rec8 cohesin
longer
thicker
compacted
F
Rec8
Rec25
Rec8-
GFP
Rec25-
mCherry
Meiotic prophase (mei4Δ wpl1Δ)
rec10Δ
enhanced

G

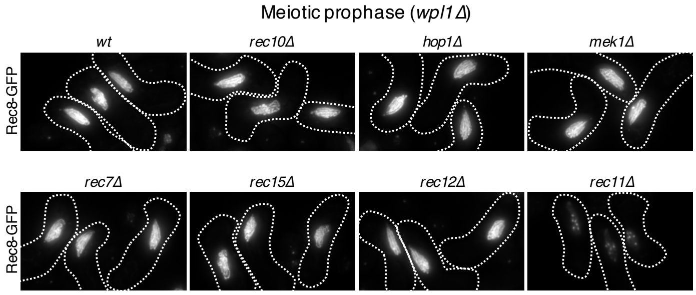


H

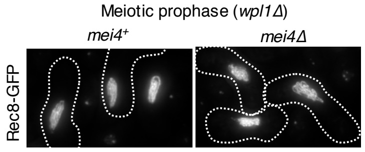


**Figure S4.** Meiotic events in the *wpl1*Δ mutant
(A) Synchronous progression of meiosis in the indicated strains in the *$h^-/h^-$ pat1-as2 MDR-sup* background monitored by the quantification of the number of nuclei stained with DAPI. (B) The degree of alignment of homologs quantified as the pairing frequency (%) of the *ade8* locus for the indicated strains. Cells were induced to meiosis at 26°C. The *ade8* loci at distance ≤0.35 µm were counted as paired, and pairing frequencies were plotted for five substages (I–V) of the horsetail stage as previously reported (61). (C) Time-lapse observation (5 min intervals) of Rec8-GFP signals with histone H2B-mCherry in the indicated strains during horsetail nuclear movement. The yellow asterisk indicates the torsional tuning of chromatin (see also Supplementary Movies 1, and 3). (D) Amplification of Hi-C contact maps for the indicated strain at VG, meiosis 0 h and 2.5 h of the 200 kbp region of chromosome 2 at 2 kbp resolution. (E) A schematic representation of meiotic chromosome organization along sister chromatids with Rec8 cohesin. In *wpl1*Δ, chromatin loops are lengthened and arrayed in certain units, resulting in a thicker Rec8 axis and a more compact chromosome. (F) Representative pictures of Rec8-GFP (green) and Rec25-mCherry (red) signals with cell shape (white contour) during meiotic prophase in wild type and *rec10*Δ in *wpl1*Δ *mei4*Δ background. An enhanced view of Rec25-mCherry in *rec10*Δ in the lower right. (G) Representative pictures of Rec8-GFP during meiotic prophase for the indicated strains in the *wpl1*Δ background, depicting cell shape (white contour). Axial Rec8-GFP signals seem to be normal in all mutants except for *rec11*Δ, which showed a great reduction in Rec8-GFP signal intensity, probably due to the loss of Rec8 cohesin from chromosomal arm regions. (H) Representative pictures of Rec8-GFP during meiotic prophase in wild-type and *mei4*Δ strains, both in a *wpl1*Δ background, depicting cell shape (white contour).

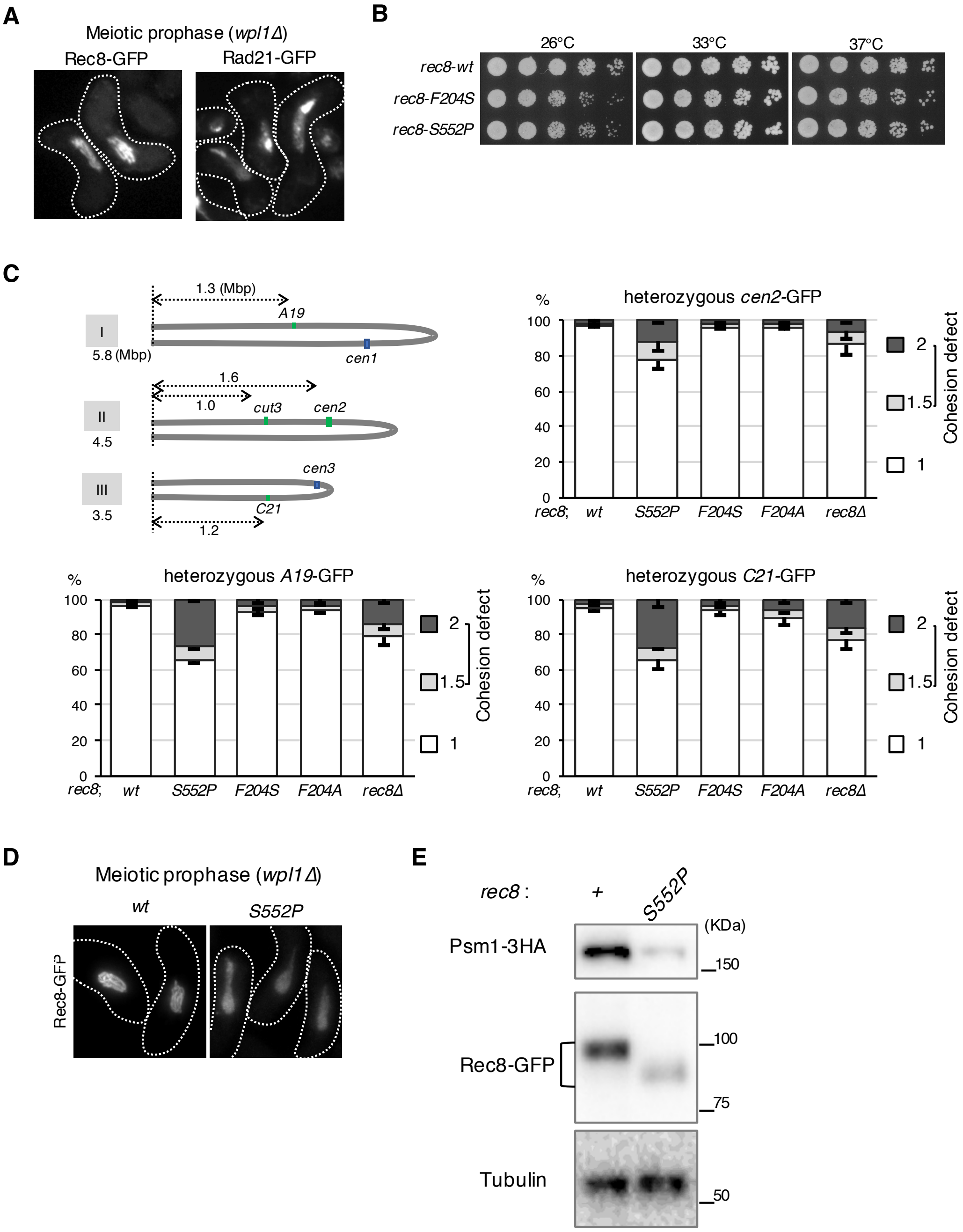
A
Meiotic prophase (wpl1Δ)
Rec8-GFP
Rad21-GFP
B
26°C
33°C
37°C
rec8-wt
rec8-F204S
rec8-S552P
C
1.3 (Mbp)
A19
I
5.8 (Mbp)
cen1
1.6
1.0
cut3
cen2
II
4.5
cen3
III
3.5
C21
1.2
heterozygous cen2-GFP
heterozygous A19-GFP
heterozygous C21-GFP
%
100
80
60
40
20
0
rec8;
wt
S552P
F204S
F204A
rec8Δ
2
1.5
1
Cohesion defect
D
Meiotic prophase (wpl1Δ)
wt
S552P
Rec8-GFP
E
rec8 :
+
S552P
(KDa)
Psm1-3HA
150
100
Rec8-GFP
75
Tubulin
50

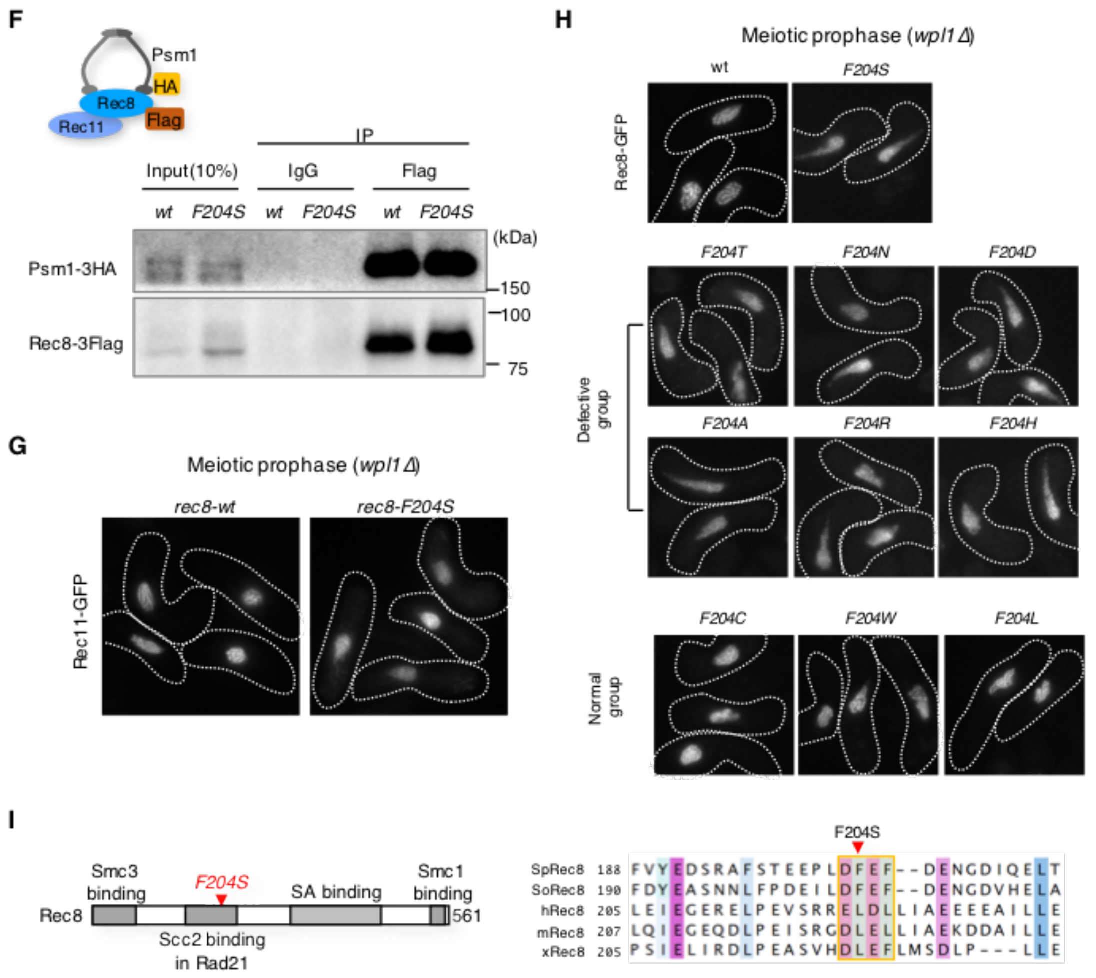


**Figure S5.** Characterization of the isolated *rec8* mutants

(A) Representative pictures of *wpl1*Δ cells during meiotic prophase. Rec8-GFP in wild type (upper) and Rad21-GFP in the *rec8*Δ *rec11*Δ strain (lower), both in the *wpl1*Δ background, with cell shape (white contour). Note that Rad21-GFP expressed in a *rec8*Δ *rec11*Δ double mutant background better mimics the mitotic cohesin complex composition (Rad21-Psc3) during meiosis, avoiding competitive actions of Rec11 (57). (B) Mitotic growth of the indicated strains expressing *rec8* under the *adh1* promoter. Five-fold serial dilutions on a YE plate at 26°C, 33°C, and 37°C. Note that although cohesion defects are expected to occur during mitosis in the *rec8-S552P* mutant, mitotic growth is ensured by their overexpression when driven by the *adh1* promoter. (C) Schematic drawing showing the positions of *lacO* insertion (*A19*, *cut3*, *cen2*, and *C21*) indicated by the distance from telomere (upper left panel). Cohesion defects at these three chromosomal loci (*A19*, *cen2*, and *C21*) were examined in the indicated strains. Cells are arrested at prophase I by the *mei4*Δ mutation. After fixation by methanol, the number of heterozygous GFP signals was counted (n > 150). A score of 2 or 1.5 of GFP signals represents the complete or partial loss of sister chromatid cohesion, respectively. Error bars show standard deviations (SD) from three independent experiments. Note that the *rec8-S552P* mutant is more deficient in cohesion than *rec8*Δ*,* probably because Rad21 can compensate for Rec8 function only in the absence of Rec8 but not in the presence of the mutant Rec8 protein. (D) Representative pictures of Rec8-GFP during meiotic prophase in wild type and *rec8-S552P* strain, both in the *wpl1*Δ background. In the *rec8-S552P*, axial Rec8-GFP signals become vague, like in the *rec8-F204S* mutant. (E) Western blot in wild type and *rec8-S552P* cells. Whole-cell extracts were prepared from meiotic prophase-arrested diploid cells expressing Psm1-3×HA with Rec8-GFP. Up-shifted Rec8-GFP signals due to the hyper phosphorylation were rarely seen in *rec8-S552P* cells, with the great reduction of the protein amount of Psm1-3×HA. This low protein expression or low stability of the Psm1 subunit in the cohesin complex in the *rec8-S552P* mutant may be responsible for the defects observed in axis formation and cohesion during meiosis (Figure S5E, 5C, and S5D). (F) Co-immunoprecipitation of Rec8-3×Flag and Psm1-3×HA. Whole-cell extracts (Input) were prepared from meiotic prophase diploid cells expressing Psm1-3×HA with Rec8-3×Flag in wild type and *rec8-F204S* cells. Rec8-3×Flag was immunoprecipitated with anti-Flag polyclonal antibodies (anti-DDDDK, MBL) to examine the co-precipitation of Psm1-3×HA. (G) Representative pictures of Rec11-GFP with cell shape (white contour) during meiotic prophase in wild type and *rec8-F204S* strain, both in *wpl1*Δ background. (H) Representative pictures of Rec8-GFP with cell shape (white contour) during meiotic prophase for the indicated *rec8* mutants in the *wpl1*Δ background. Although rec8-*F204S, F204T, F204N, F204D, F204A, F204*R and *F204H* showed defect in the formation of Rec8-GFP axis, the substitution for hydrophobic resides (*rec8-F204C*, *F204W*, and *F204L*) showed relatively normal formation of Rec8-GFP axis structures. (I) The left panel is a schematic diagram of *S. pombe* Rec8. Domains for the known interaction with cohesin-related factors are shown. The position of F204 is also indicated by the red arrowhead. The right panel depicts the alignment of Rec8 homologs in *Shizosaccharomyces pombe* (Sp), *S. octosporus* (So), human (h), mouse (m) and frog (*Xenopus leaves*) (x).

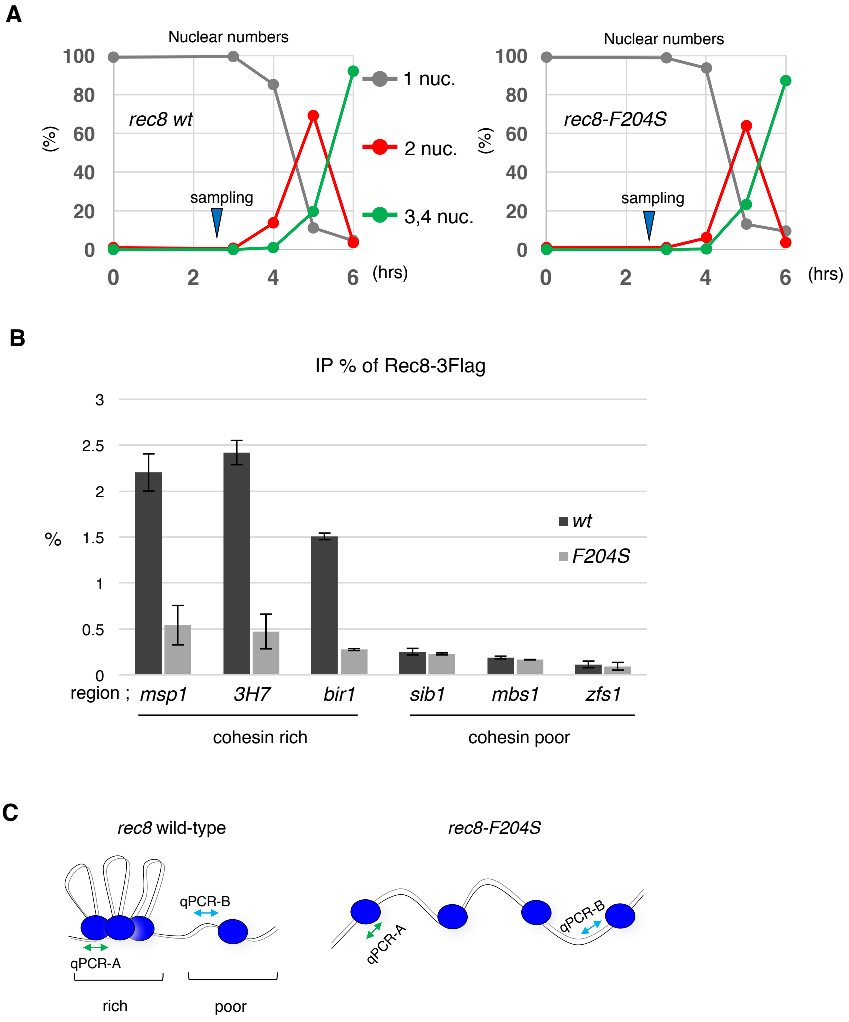


**Figure S6. Defective chromatin loop formation in the *rec8-F204S* mutant**
(A) Synchronous progression of meiosis in the indicated strains in the *h⁻/h⁻ pat1-as2 MDR-sup* background, monitored by the quantification of the number of nuclei stained by DAPI. (B) Chromatin immunoprecipitation assay for Rec8 binding. Rec8 binding at the chromosome arm regions [*msp1, SPBC3H7.03(3H7), bir1, sib1, mbs1,* and *zfs1*] were examined using ChIP assay with anti-Flag-M2 antibody and mouse non-immune IgG. Each percent IP of mouse non-immune IgG were used to subtract from that of Flag-M2 antibody. Each percent IP was quantified from three independent qPCR experiments. Error bars show SD. (C) A diagram illustrating the loss of loop structure in the *rec8-F204S* mutant strain, which results in a reduced percent IP in qPCR in the cohesin-rich region by ChIP. In the case of the qPCR-A region in this example, the *rec8* wild type is about three times more likely to co-precipitate with Rec8 than *rec8-F204S* mutant, and qPCR-B is unchanged between the two strains.

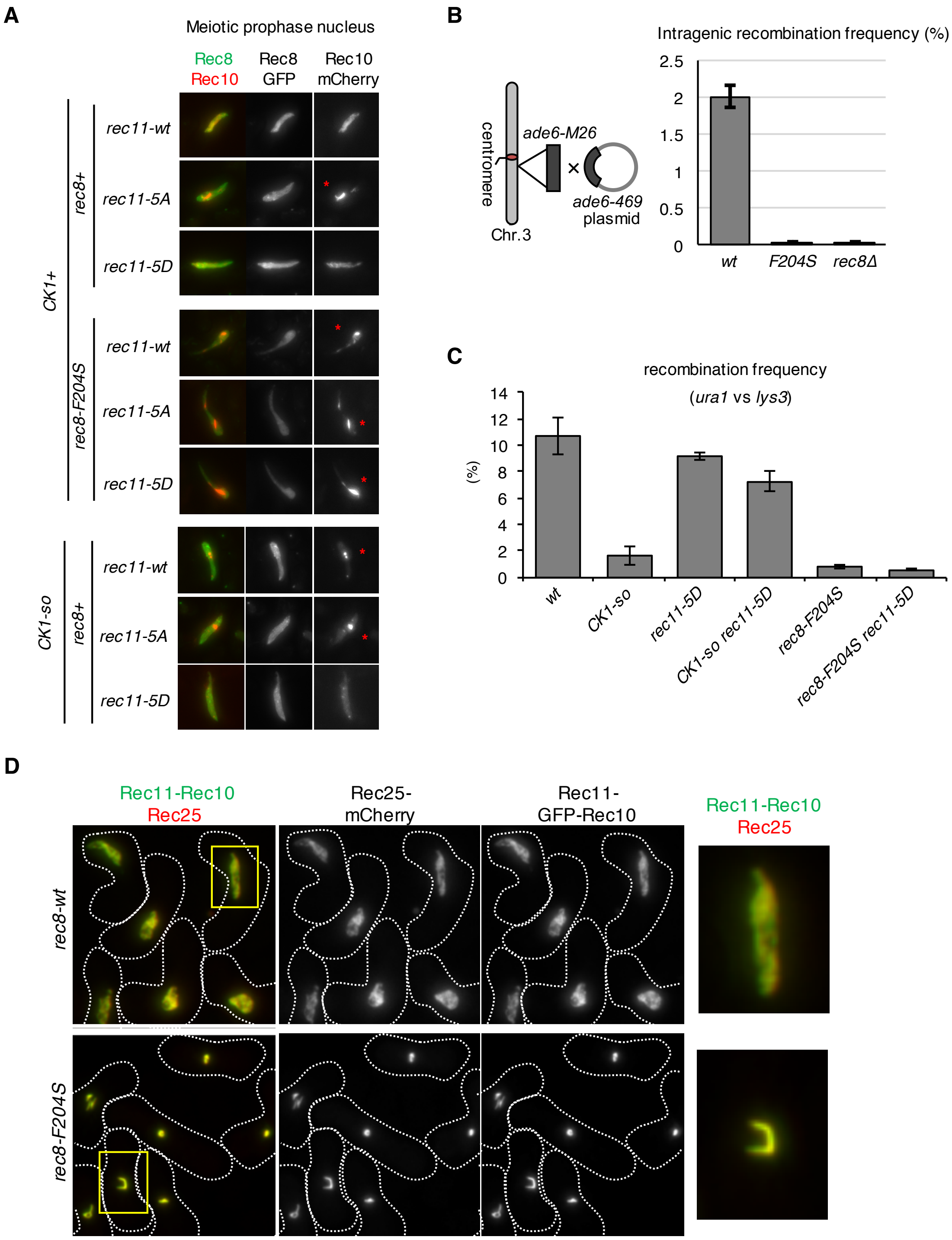


**Figure S7.** Roles for Rec8 in LinE assembly and meiotic recombination
(A) Representative pictures of Rec8-GFP (green) and Rec10-mCherry (red) during meiotic prophase. A wild type or mutant *rec11* construct was expressed in the indicated strains. Red asterisks indicate Rec10 aggregates. In case of the CK1 wild type background, *rec11-5A* which lost the CK1-dependent phosphorylation, showed defects in LinE assembly (Rec10 forms aggregates), but *rec11-5D* shows normal LinE formation. Although introduction of the *rec11-5D* allele can suppress the LinE formation

defect in the *CK1-so* mutant, but not in the *rec8-F204S* mutant, suggesting that LinE factors such as Rec10 cannot localize to chromatin stably, even though the CK1-dependent Rec8-Rec11 cohesin recruitment pathway functions normally in *rec8-F204S* mutant, as shown in Figure 7B. Thus, CK1-dependent recruitment pathway of LinE factors via Rec8-Rec11 cohesin to chromatin can no longer function in the *rec8-F204S* mutant strain. (B) Intragenic recombination frequency at the *ade6* locus, between *ade6-M26* and *ade6-469*. Error bars show standard deviations (SD) from three independent experiment (n>7400, each). (C) Intergenic meiotic recombination in the *ura1-lys3* (n > 300) assessed in the indicated zygotes. Error bars show standard deviations (SD) from three independent experiment. (D) Representative pictures of Rec11-GFP-Rec10 fusion protein (green) and Rec25-mCherry (red) with cell shape (white contour) during meiotic prophase for the indicated strains. A previous study demonstrated that the expression of Rec11-GFP-Rec10 fusion protein can suppress the LinE formation defect caused by the loss of CK1-dependet phosphorylation of Rec11 (48). However, the expression of Rec11-GFP-Rec10 fusion protein in *rec8-F204S* mutant formed Rec25 aggregates together with Rec11-GFP-Rec10 fusion protein itself. This implies that even if Rec10 is once recruited to chromatin by fusion with Rec11, the loss of the Rec8-dependent axis-loop chromatin structure in the *rec8-F204S* mutant does not stabilize its localization on chromatin. This eventually leads to the aggregation with other LinE factors (such as Rec25, etc.), probably because its collective force of LinE aggregates is stronger than the binding force of Rec8 and Rec11.

**Supplementary Table S1.** List of *S. pombe* strains used in this study.

| | | |
|---|---|---|
| **Figure 1B, C and D** | | |
| YS193 | *h-/h-* | *ade6-M210/M216 leu1/leu1 ura4-D18/ura4-D18 pat1-L95A<<bleo/pat1-L95A<<bleo caf5::bsdr/caf5::bsdr pap1-del/pap1-del pmd1-del/pmd1-del mfs1-del/mfs1-del bfr1-del/bfr1-del dnf2-del/dnf2-del erg5::ura4$^{+}$/erg5::ura4$^{+}$ +/CO2::mat-Pc-natr rec8$^{+}$-GFP-kanr/rec8$^{+}$-GFP-kanr* |
| **Figure 2A and B** | | |
| YS753 | *h-/h-* | *ade6-M216/210 leu1/leu1 ura4-D18/ura4-D18 pat1-L95A<<bleo/pat1-L95A<<bleo caf5::bsdr/caf5::bsdr pap1-del/pap1-del pmd1-del/pmd1-del mfs1-del/mfs1-del bfr1-del/bfr1-del dnf2-del/dnf2-del erg5::ura4$^{+}$/erg5::ura4$^{+}$ +/CO2::mat-Pc-natr rec8$^{+}$-GFP-kanr/rec8$^{+}$-GFP-kanr Δrec10::hygr/Δrec10::hygr* |
| YS250 | *h-/h-* | *ade6-M210/M216 leu1/leu1 ura4-D18/ura4-D18 pat1-L95A<<bleo/pat1-L95A<<bleo caf5::bsdr/caf5::bsdr pap1-del/pap1-del pmd1-del/pmd1-del mfs1-del/mfs1-del bfr1-del/bfr1-del dnf2-del/dnf2-del erg5::ura4$^{+}$/erg5::ura4$^{+}$ +/CO2::mat-Pc-natr rec8$^{+}$-GFP-kanr/rec8$^{+}$-GFP-kanr Δrec12::hygr/Δrec12::hygr* |
| YS249 | *h-/h-* | *ade6-M210/M216 leu1/leu1 ura4-D18/ura4-D18 pat1-L95A<<bleo/pat1-L95A<<bleo caf5::bsdr/caf5::bsdr pap1-del/pap1-del pmd1-del/pmd1-del mfs1-del/mfs1-del bfr1-del/bfr1-del dnf2-del/dnf2-del erg5::ura4$^{+}$/erg5::ura4$^{+}$ +/CO2::mat-Pc-natr Δrec8::kanr/Δrec8::kanr* |
| **Figure 3A and D** | | |
| YS193 | *h-/h-* | *ade6-M210/M216 leu1/leu1 ura4-D18/ura4-D18 pat1-L95A<<bleo/pat1-L95A<<bleo caf5::bsdr/caf5::bsdr pap1-del/pap1-del pmd1-del/pmd1-del mfs1-del/mfs1-del bfr1-del/bfr1-del dnf2-del/dnf2-del erg5::ura4$^{+}$/erg5::ura4$^{+}$ +/CO2::mat-Pc-natr rec8$^{+}$-GFP-kanr/rec8$^{+}$-GFP-kanr* |
| YS753 | *h-/h-* | *ade6-M216/210 leu1/leu1 ura4-D18/ura4-D18 pat1-L95A<<bleo/pat1-L95A<<bleo caf5::bsdr/caf5::bsdr pap1-del/pap1-del pmd1-del/pmd1-del mfs1-del/mfs1-del bfr1-del/bfr1-del dnf2-del/dnf2-del erg5::ura4$^{+}$/erg5::ura4$^{+}$ +/CO2::mat-Pc-natr rec8$^{+}$-GFP-kanr/rec8$^{+}$-GFP-kanr Δrec10::hygr/Δrec10::hygr* |
| YS250 | *h-/h-* | *ade6-M210/M216 leu1/leu1 ura4-D18/ura4-D18 pat1-L95A<<bleo/pat1-L95A<<bleo caf5::bsdr/caf5::bsdr pap1-del/pap1-del pmd1-del/pmd1-del mfs1-del/mfs1-del bfr1-del/bfr1-del dnf2-del/dnf2-del erg5::ura4$^{+}$/erg5::ura4$^{+}$ +/CO2::mat-Pc-natr rec8$^{+}$-GFP-kanr/rec8$^{+}$-GFP-kanr Δrec12::hygr/Δrec12::hygr* |
| YS249 | *h-/h-* | *ade6-M210/M216 leu1/leu1 ura4-D18/ura4-D18 pat1-L95A<<bleo/pat1-L95A<<bleo caf5::bsdr/caf5::bsdr pap1-del/pap1-del pmd1-del/pmd1-del mfs1-del/mfs1-del bfr1-del/bfr1-del dnf2-del/dnf2-del erg5::ura4$^{+}$/erg5::ura4$^{+}$ +/CO2::mat-Pc-natr Δrec8::kanr/Δrec8::kanr* |
| **Figure 3B, C and E** | | |
| YS193 | *h-/h-* | *ade6-M210/M216 leu1/leu1 ura4-D18/ura4-D18 pat1-L95A<<bleo/pat1-L95A<<bleo caf5::bsdr/caf5::bsdr pap1-del/pap1-del pmd1-del/pmd1-del mfs1-del/mfs1-del bfr1-del/bfr1-del dnf2-del/dnf2-del erg5::ura4$^{+}$/erg5::ura4$^{+}$ +/CO2::mat-Pc-natr rec8$^{+}$-GFP-kanr/rec8$^{+}$-GFP-kanr* |
| **Figure 4A** | | |
| PY249 | *h90* | *Δmei4::ura4$^{+}$ rec8$^{+}$-GFP-kanr* |
| PC345 | *h90* | *Δmei4::ura4$^{+}$ rec8$^{+}$-GFP-kanr Δwpl1::hygr* |
| **Figure 4B, C, D and E** | | |
| YS193 | *h-/h-* | *ade6-M210/M216 leu1/leu1 ura4-D18/ura4-D18 pat1-L95A<<bleo/pat1-L95A<<bleo caf5::bsdr/caf5::bsdr pap1-del/pap1-del pmd1-del/pmd1-del mfs1-del/mfs1-del bfr1-del/bfr1-del dnf2-del/dnf2-del erg5::ura4$^{+}$/erg5::ura4$^{+}$ +/CO2::mat-Pc-natr rec8$^{+}$-GFP-kanr/rec8$^{+}$-GFP-kanr* |
| YS248 | *h-/h-* | *ade6-M210/M216 leu1/leu1 ura4-D18/ura4-D18 pat1-L95A<<bleo/pat1-L95A<<bleo caf5::bsdr/caf5::bsdr pap1-del/pap1-del pmd1-del/pmd1-del mfs1-del/mfs1-del bfr1-del/bfr1-del dnf2-del/dnf2-del erg5::ura4$^{+}$/erg5::ura4$^{+}$ +/CO2::mat-Pc-natr rec8$^{+}$-GFP-kanr/rec8$^{+}$-GFP-kanr Δwpl1::hygr/Δwpl1::hygr* |
| **Figure 5B** | | |
| YS036 | *h90* | *Δmei4::ura4$^{+}$ Δpsc3::kanr ura4-Padh41-rec11Δwpl1::hygr Δrad21::ura4$^{+}$ ade6? Padh1-rec8$^{+}$-GFP-bsdr* |

| | | |
|---|---|---|
| YS038 | *h90* | *Δmei4::ura4$^+$ Δpsc3::kanr ura4-Padh41-rec11Δwpl1::hygr Δrad21::ura4$^+$ ade6? Padh1-rec8-F204S-GFP-bsdr* |
| **Figure 5C** | | |
| YS519 | *h+* | *rec8$^+$-3Pk-bsdr ade6-M216 leu1 Δmei4::hygr* |
| YS635 | *h-* | *cut3-lacOp his7$^+$::lacI-GFP rec8$^+$-3Pk-bsdr Δmei4::ura4$^+$* |
| YS520 | *h+* | *rec8-S552P-3Pk-bsdr ade6-M216 leu1 Δmei4::hygr* |
| YS638 | *h-* | *cut3-lacOp his7$^+$::lacI-GFP rec8-S552P-3Pk-bsdr Δmei4::ura4$^+$* |
| YS522 | *h+* | *rec8-F204S-3Pk-bsdr leu1 Δmei4::hygr* |
| YS641 | *h-* | *cut3-lacOp his7$^+$::lacI-GFP rec8-F204S-3Pk-bsdr Δmei4::ura4$^+$* |
| YS562 | *h+* | *Δmei4::ura4$^+$ ura4 ade6-M210 leu1 Δrec8::kanr* |
| YS646 | *h-* | *cut3-lacOp his7$^+$::lacI-GFP Δrec8::kanr Δmei4::ura4$^+$ leu1* |
| **Figure 5D** | | |
| YS686 | *h90* | *rec8$^+$-GFP-bsdr ura4$^+$ ade6? aur1$^+$::htb1-mCherry* |
| YS688 | *h90* | *rec8-F204S-GFP-bsdr ura4$^+$ ade6? aur1$^+$::htb1-mCherry* |
| YS689 | *h90* | *rec8::kanr ade6 aur1$^+$::htb1-mCherry* |
| **Figure 5E** | | |
| YS692 | *h-* | *rec8$^+$-3Pk-bsdr lys1$^+$-taz1-GFP ade8-lacO-kanr-GFP leu1* |
| YS485 | *h+* | *rec8$^+$-3Pk-bsdr ade6-M216 leu1* |
| YS693 | *h-* | *rec8-F204S-3Pk-bsdr lys1$^+$-taz1-GFP ade8-lacO-kanr-GFP leu1* |
| YS420 | *h+* | *rec8-F204S-3Pk-bsdr leu1* |
| YS695 | *h-* | *Δrec8::kanr lys1$^+$-taz1-GFP ade8-lacO-kanr-GFP leu1* |
| YS561 | *h+* | *ade6-M216 leu1 Δrec8::kanr* |
| **Figure 6A, B, C and D** | | |
| YS547 | *h-/h-* | *ade6-M216/210 leu1/leu1 ura4-D18/ura4-D18 pat1-L95A<<bleo/pat1-L95A<<bleo caf5::bsdr/caf5::bsdr pap1-del/pap1-del pmd1-del/pmd1-del mfs1-del/mfs1-del bfr1-del/bfr1-del dnf2-del/dnf2-del erg5::ura4$^+$/erg5::ura4$^+$ +/CO2::mat-Pc-natr rec8$^+$-3Flag-kanr/rec8$^+$-3Flag-kanr* |
| YS549 | *h-/h-* | *ade6-M216/210 leu1/leu1 ura4-D18/ura4-D18 pat1-L95A<<bleo/pat1-L95A<<bleo caf5::bsdr/caf5::bsdr pap1-del/pap1-del pmd1-del/pmd1-del mfs1-del/mfs1-del bfr1-del/bfr1-del dnf2-del/dnf2-del erg5::ura4$^+$/erg5::ura4$^+$ +/CO2::mat-Pc-natr rec8-F204S-3Flag-kanr/rec8-F204S-3Flag-kanr* |
| **Figure 6E** | | |
| YS547 | *h-/h-* | *ade6-M216/210 leu1/leu1 ura4-D18/ura4-D18 pat1-L95A<<bleo/pat1-L95A<<bleo caf5::bsdr/caf5::bsdr pap1-del/pap1-del pmd1-del/pmd1-del mfs1-del/mfs1-del bfr1-del/bfr1-del dnf2-del/dnf2-del erg5::ura4$^+$/erg5::ura4$^+$ +/CO2::mat-Pc-natr rec8$^+$-3Flag-kanr/rec8$^+$-3Flag-kanr* |
| YS549 | *h-/h-* | *ade6-M216/210 leu1/leu1 ura4-D18/ura4-D18 pat1-L95A<<bleo/pat1-L95A<<bleo caf5::bsdr/caf5::bsdr pap1-del/pap1-del pmd1-del/pmd1-del mfs1-del/mfs1-del bfr1-del/bfr1-del dnf2-del/dnf2-del erg5::ura4$^+$/erg5::ura4$^+$ +/CO2::mat-Pc-natr rec8-F204S-3Flag-kanr/rec8-F204S-3Flag-kanr* |
| YS249 | *h-/h-* | *ade6-M210/M216 leu1/leu1 ura4-D18/ura4-D18 pat1-L95A<<bleo/pat1-L95A<<bleo caf5::bsdr/caf5::bsdr pap1-del/pap1-del pmd1-del/pmd1-del mfs1-del/mfs1-del bfr1-del/bfr1-del dnf2-del/dnf2-del erg5::ura4$^+$/erg5::ura4$^+$ +/CO2::mat-Pc-natr Δrec8::kanr/Δrec8::kanr* |
| **Figure 7A** | | |
| PB274 | *h90* | *cut3-lacOp his7$^+$::lacI-GFP mei4::ura4$^+$ leu1 rec10$^+$-mCherry-natr* |
| YS387 | *h90* | *cut3-lacOp his7$^+$::lacI-GFP mei4::ura4$^+$ leu1 rec10$^+$-mCherry-natr rec8-F204S-3Pk-bsdr* |
| YS132 | *h90* | *cut3-lacOp his7$^+$::lacI-GFP mei4::ura4$^+$ leu1 rec10$^+$-mCherry-natr Δrec8::kanr* |
| **Figure 7B** | | |
| YS1278 | *h+* | *rec8$^+$-3Pk-bsdr ade6-M216 leu1 vps38$^+$ -1000::natr* |
| YS1282 | *h-* | *ura1 lys3 rec8$^+$-3Pk-bsdr cox7$^+$ -1215::hygr* |
| YS1279 | *h+* | *rec8-F204S-3Pk-bsdr leu1 vps38$^+$ -1000::natr* |
| YS1283 | *h-* | *ura1 lys3 rec8-F204S-3Pk-bsdr cox7$^+$ -1215::hygr* |
| YS1280 | *h+* | *ade6-M216 leu1 Δrec8::kanr vps38$^+$ -1000::natr* |
| YS1284 | *h-* | *ura1 lys3 Δrec8::kanr cox7$^+$ -1215::hygr* |

| | | |
|---|---|---|
| **Figure 7C** | | |
| YS705 | *h-/h-* | *ade6-M216/210 leu1/leu1 ura4-D18/ura4-D18 pat1-L95A<<bleo/pat1-L95A<<bleo caf5::bsdr/caf5::bsdr pap1-del/pap1-del pmd1-del/pmd1-del mfs1-del/mfs1-del bfr1-del/bfr1-del dnf2-del/dnf2-del erg5::ura4$^+$/erg5::ura4$^+$ +/CO2::mat-Pc-natr rec8$^+$-3Flag-kanr/rec8$^+$-3Flag-kanr psm1-3HA-hygr/psm1-3HA-hygr* |
| YS706 | *h-/h-* | *ade6-M216/210 leu1/leu1 ura4-D18/ura4-D18 pat1-L95A<<bleo/pat1-L95A<<bleo caf5::bsdr/caf5::bsdr pap1-del/pap1-del pmd1-del/pmd1-del mfs1-del/mfs1-del bfr1-del/bfr1-del dnf2-del/dnf2-del erg5::ura4$^+$/erg5::ura4$^+$ +/CO2::mat-Pc-natr rec8-F204S-3Flag-kanr/rec8-F204S-3Flag-kanr psm1-3HA-hygr/psm1-3HA-hygr* |
| **Figure S1A, B and D** | | |
| YS193 | *h-/h-* | *ade6-M210/M216 leu1/leu1 ura4-D18/ura4-D18 pat1-L95A<<bleo/pat1-L95A<<bleo caf5::bsdr/caf5::bsdr pap1-del/pap1-del pmd1-del/pmd1-del mfs1-del/mfs1-del bfr1-del/bfr1-del dnf2-del/dnf2-del erg5::ura4$^+$/erg5::ura4$^+$ +/CO2::mat-Pc-natr rec8$^+$-GFP-kanr/rec8$^+$-GFP-kanr* |
| **Figure S1C** | | |
| YS1106 | *h-/h-* | *ade6-M210/ade6-M216 leu1/leu1 ura4-D18/ura4-D18 pat1-L95A<<bleo/pat1-L95A<<bleo caf5::bsdr/caf5::bsdr pap1-del /pap1-del pmd1-del/pmd1-del mfs1-del/mfs1-del bfr1-del /bfr1-del dnf2-del/dnf2-del erg5::ura4$^+$ /erg5::ura4$^+$ rec8-3flag-kanr/rec8-3flag-kanr mis4-13myc-hygr/mis4-13myc-hygr +/CO2::mat-Pc-natr* |
| **Figure S2** | | |
| YS753 | *h-/h-* | *ade6-M216/210 leu1/leu1 ura4-D18/ura4-D18 pat1-L95A<<bleo/pat1-L95A<<bleo caf5::bsdr/caf5::bsdr pap1-del/pap1-del pmd1-del/pmd1-del mfs1-del/mfs1-del bfr1-del/bfr1-del dnf2-del/dnf2-del erg5::ura4$^+$/erg5::ura4$^+$ +/CO2::mat-Pc-natr rec8$^+$-GFP-kanr/rec8$^+$-GFP-kanr Δrec10::hygr/Δrec10::hygr* |
| YS250 | *h-/h-* | *ade6-M210/M216 leu1/leu1 ura4-D18/ura4-D18 pat1-L95A<<bleo/pat1-L95A<<bleo caf5::bsdr/caf5::bsdr pap1-del/pap1-del pmd1-del/pmd1-del mfs1-del/mfs1-del bfr1-del/bfr1-del dnf2-del/dnf2-del erg5::ura4$^+$/erg5::ura4$^+$ +/CO2::mat-Pc-natr rec8$^+$-GFP-kanr/rec8$^+$-GFP-kanr Δrec12::hygr/Δrec12::hygr* |
| YS249 | *h-/h-* | *ade6-M210/M216 leu1/leu1 ura4-D18/ura4-D18 pat1-L95A<<bleo/pat1-L95A<<bleo caf5::bsdr/caf5::bsdr pap1-del/pap1-del pmd1-del/pmd1-del mfs1-del/mfs1-del bfr1-del/bfr1-del dnf2-del/dnf2-del erg5::ura4$^+$/erg5::ura4$^+$ +/CO2::mat-Pc-natr Δrec8::kanr/Δrec8::kanr* |
| **Figure S3A** | | |
| YS193 | *h-/h-* | *ade6-M210/M216 leu1/leu1 ura4-D18/ura4-D18 pat1-L95A<<bleo/pat1-L95A<<bleo caf5::bsdr/caf5::bsdr pap1-del/pap1-del pmd1-del/pmd1-del mfs1-del/mfs1-del bfr1-del/bfr1-del dnf2-del/dnf2-del erg5::ura4$^+$/erg5::ura4$^+$ +/CO2::mat-Pc-natr rec8$^+$-GFP-kanr/rec8$^+$-GFP-kanr* |
| YS753 | *h-/h-* | *ade6-M216/210 leu1/leu1 ura4-D18/ura4-D18 pat1-L95A<<bleo/pat1-L95A<<bleo caf5::bsdr/caf5::bsdr pap1-del/pap1-del pmd1-del/pmd1-del mfs1-del/mfs1-del bfr1-del/bfr1-del dnf2-del/dnf2-del erg5::ura4$^+$/erg5::ura4$^+$ +/CO2::mat-Pc-natr rec8$^+$-GFP-kanr/rec8$^+$-GFP-kanr Δrec10::hygr/Δrec10::hygr* |
| YS250 | *h-/h-* | *ade6-M210/M216 leu1/leu1 ura4-D18/ura4-D18 pat1-L95A<<bleo/pat1-L95A<<bleo caf5::bsdr/caf5::bsdr pap1-del/pap1-del pmd1-del/pmd1-del mfs1-del/mfs1-del bfr1-del/bfr1-del dnf2-del/dnf2-del erg5::ura4$^+$/erg5::ura4$^+$ +/CO2::mat-Pc-natr rec8$^+$-GFP-kanr/rec8$^+$-GFP-kanr Δrec12::hygr/Δrec12::hygr* |
| YS249 | *h-/h-* | *ade6-M210/M216 leu1/leu1 ura4-D18/ura4-D18 pat1-L95A<<bleo/pat1-L95A<<bleo caf5::bsdr/caf5::bsdr pap1-del/pap1-del pmd1-del/pmd1-del mfs1-del/mfs1-del bfr1-del/bfr1-del dnf2-del/dnf2-del erg5::ura4$^+$/erg5::ura4$^+$ +/CO2::mat-Pc-natr Δrec8::kanr/Δrec8::kanr* |
| **Figure S3B** | | |
| YS193 | *h-/h-* | *ade6-M210/M216 leu1/leu1 ura4-D18/ura4-D18 pat1-L95A<<bleo/pat1-L95A<<bleo caf5::bsdr/caf5::bsdr pap1-del/pap1-del pmd1-del/pmd1-del mfs1-del/mfs1-del bfr1-del/bfr1-del dnf2-del/dnf2-del erg5::ura4$^+$/erg5::ura4$^+$ +/CO2::mat-Pc-natr rec8$^+$-GFP-kanr/rec8$^+$-GFP-kanr* |
| **Figure S4A and D** | | |
| YS193 | *h-/h-* | *ade6-M210/M216 leu1/leu1 ura4-D18/ura4-D18 pat1-L95A<<bleo/pat1-L95A<<bleo caf5::bsdr/caf5::bsdr pap1-del/pap1-del pmd1-del/pmd1-del mfs1-del/mfs1-del bfr1-del/bfr1-del dnf2-del/dnf2-del erg5::ura4$^+$/erg5::ura4$^+$ +/CO2::mat-Pc-natr rec8$^+$-GFP-kanr/rec8$^+$-GFP-kanr* |

| | | |
|---|---|---|
| YS248 | *h-/h-* | *ade6-M210/M216 leu1/leu1 ura4-D18/ura4-D18 pat1-L95A<<bleo/pat1-L95A<<bleo caf5::bsdr/caf5::bsdr pap1-del/pap1-del pmd1-del/pmd1-del mfs1-del/mfs1-del bfr1-del/bfr1-del dnf2-del/dnf2-del erg5::ura4$^{+}$/erg5::ura4$^{+}$ +/CO2::mat-Pc-natr rec8$^{+}$-GFP-kanr/rec8$^{+}$-GFP-kanr Δwpl1::hygr/Δwpl1::hygr* |
| **Figure S4B** | | |
| YW295-26C | *h90* | *ade6-216 leu1 ura4 ade8-kanr-ura4$^{+}$-lacOp his7$^{+}$::lacI-GFP* |
| YY313-1B | *h-* | *leu1 ura4 ade6? lys1? Δpds5::LEU2 ade8-kanr-ura4-lacOp his7$^{+}$::lacI-GFP* |
| YY313-4D | *h+* | *leu1 ura4 ade6? lys1? Δpds5::LEU2 ade8-kanr-ura4-lacOp his7$^{+}$::lacI-GFP* |
| MK104 | *h-* | *leu1 lys1 ura4 Δrec8::ura4$^{+}$ade8-kanr-ura4-lacOp his7$^{+}$::lacI-GFP* |
| AY266-5A | *h+* | *lys1 ura4 Δrec8::ura4$^{+}$ ade8-kanr-ura4-lacOp his7$^{+}$::lacI-GFP* |
| YY760-7C | *h90* | *ura4 leu1 lys1 Δwpl1::hygr ade8-kanr-ura4-lacOp his7$^{+}$::lacI-GFP* |
| **Figure S4C** | | |
| YS686 | *h90* | *rec8$^{+}$-GFP-bsdr ade6? aur1$^{+}$::htb1-mCherry* |
| YS1152 | *h90* | *rec8$^{+}$-GFP-bsdr ade6? aur1$^{+}$::htb1-mCherry Δwpl1::natr* |
| **Figure S4F** | | |
| YS1175 | *h90* | *Δmei4::ura4$^{+}$ rec8$^{+}$-GFP-kanr Δwpl1::hygr rec25-mCherry-natr* |
| YS1176 | *h90* | *Δmei4::ura4$^{+}$ rec8$^{+}$-GFP-kanr Δwpl1::hygr rec10::bsdr ade6? rec25-mCherry-natr* |
| **Figure S4G** | | |
| PC345 | *h90* | *Δmei4::ura4$^{+}$ rec8$^{+}$-GFP-kanr Δwpl1::hygr* |
| YS1154 | *h90* | *Δmei4::ura4$^{+}$ rec8$^{+}$-GFP-kanr Δwpl1::hygr Δrec10::bsdr (ade6)* |
| YS772 | *h90* | *Δmei4::ura4$^{+}$ rec8$^{+}$-GFP-kanr Δwpl1::hygr Δhop1::bsdr* |
| YS814 | *h90* | *Δmei4::ura4$^{+}$ rec8$^{+}$-GFP-kanr Δwpl1::hygr Δmek1::natr* |
| YS1100 | *h90* | *Δmei4::ura4$^{+}$ rec8$^{+}$-GFP-kanr Δwpl1::hygr Δrec7::bsdr* |
| YS1102 | *h90* | *Δmei4::ura4$^{+}$ rec8$^{+}$-GFP-kanr Δwpl1::hygr Δrec15::bsdr* |
| YS1272 | *h90* | *Δmei4::ura4$^{+}$ rec8$^{+}$-GFP-kanr Δwpl1::hygr Δrec12::bsdr* |
| YS223 | *h90* | *Δmei4::ura4$^{+}$ Δrec11::kanr Δwpl1::hyg.B rec8$^{+}$-GFP-kanr ade6-M216* |
| **Figure S4H** | | |
| PC345 | *h90* | *Δmei4::ura4$^{+}$ rec8$^{+}$-GFP-kanr Δwpl1::hygr* |
| YS172 | *h90* | *ade6-M216 rec8$^{+}$-GFP-kanr Δwpl1::hygr* |
| **Figure S5A** | | |
| PC345 | *h90* | *Δmei4::ura4$^{+}$ rec8$^{+}$-GFP-kanr Δwpl1::hygr* |
| YS760 | *h90* | *ade6-M216? Δrec8::Padh1-rad21-GFP-ura4$^{+}$ Δrad21::kanr Δrec11::natr Δwpl1::hygr* |
| **Figure S5B** | | |
| YS036 | *h90* | *Δmei4::ura4$^{+}$ Δpsc3::kanr ura4-Padh41-rec11Δwpl1::hygr Δrad21::ura4$^{+}$ ade6? Padh1-rec8$^{+}$-GFP-bsdr* |
| YS038 | *h90* | *Δmei4::ura4$^{+}$ Δpsc3::kanr ura4-Padh41-rec11Δwpl1::hygr Δrad21::ura4$^{+}$ ade6? Padh1-rec8-F204S-GFP-bsdr* |
| YS037 | *h90* | *Δmei4::ura4$^{+}$ Δpsc3::kanr ura4-Padh41-rec11Δwpl1::hygr Δrad21::ura4$^{+}$ ade6? Padh1-rec8-S552P-GFP-bsdr* |
| **Figure S5C** | | |
| YS519 | *h+* | *rec8$^{+}$-3Pk-bsdr ade6-M216 leu1 Δmei4::hygr* |
| YS520 | *h+* | *rec8-S552P-3Pk-bsdr ade6-M216 leu1 Δmei4::hygr* |
| YS522 | *h+* | *rec8-F204S-3Pk-bsdr leu1 Δmei4::hygr* |
| YS523 | *h+* | *rec8-F204A-3Pk-bsdr leu1 Δmei4::hygr* |
| YS562 | *h+* | *Δmei4::ura4$^{+}$ ura4 ade6-M210 leu1 Δrec8::kanr* |
| YS664 | *h-* | *A19-kanr-ura4-lacO his7$^{+}$::lacI-GFP Δmei4::natr rec8$^{+}$-3Pk-bsdr ade6-M216?* |
| YS665 | *h-* | *A19-kanr-ura4-lacO his7$^{+}$::lacI-GFP Δmei4::natr rec8-S552P-3Pk-bsdr ade6-M216?* |
| YS666 | *h-* | *A19-kanr-ura4-lacO his7$^{+}$::lacI-GFP Δmei4::natr rec8-F204S-3Pk-bsdr* |
| YS667 | *h-* | *A19-kanr-ura4-lacO his7$^{+}$::lacI-GFP Δmei4::natr rec8-F204A-3Pk-bsdr* |
| YS668 | *h-* | *A19-kanr-ura4-lacO his7$^{+}$::lacI-GFP Δmei4::natr Δrec8::kanr* |
| YS679 | *h-* | *C21-kanr-ura4-lacO his7$^{+}$::lacI-GFP Δmei4::natr rec8$^{+}$-3Pk-bsdr ade6-M216?* |
| YS680 | *h-* | *C21-kanr-ura4-lacO his7$^{+}$::lacI-GFP Δmei4::natr rec8-S552P-3Pk-bsdr ade6-M216?* |
| YS681 | *h-* | *C21-kanr-ura4-lacO his7$^{+}$::lacI-GFP Δmei4::natr rec8-F204S-3Pk-bsdr* |

| | | |
|---|---|---|
| YS682 | h- | C21-kanr-ura4-lacO his7$^{+}$::lacI-GFP Δmei4::natr rec8-F204A-3Pk-bsdr |
| YS683 | h- | C21-kanr-ura4-lacO his7$^{+}$::lacI-GFP Δmei4::natr Δrec8::kanr |
| YS600 | h- | cen2-kanr-ura4-lacO his7$^{+}$::lacI-GFP rec8$^{+}$-3Pk-bsdr leu1 pnatrZA13-mCherry-atb2 Δmei4::hygr |
| YS601 | h- | cen2-kanr-ura4-lacO his7$^{+}$::lacI-GFPrec8-F204S-3Pk-bsdr leu1 pnatrZA13-mCherry-atb2 Δmei4::hygr |
| YS602 | h- | cen2-kanr-ura4-lacO his7$^{+}$::lacI-GFPrec8-F204A-3Pk-bsdr leu1 pnatrZA13-mCherry-atb2 Δmei4::hygr |
| YS603 | h- | cen2-kanr-ura4-lacO his7$^{+}$::lacI-GFP leu1 Δrec12::LEU2 z::natr-Padh13-mCherry-atb2 ade6 Δmei4::hygr |
| YS609 | h- | cen2-kanr-ura4-lacO his7$^{+}$::lacI-GFPleu1 Δrec12::LEU2 z::natr-Padh13-mCherry-atb2 ade6 Δmei4::hygr Δrec8::bsdr |
| **Figure S5D** | | |
| PC345 | h90 | Δmei4::ura4$^{+}$ rec8$^{+}$-GFP-kanr Δwpl1::hygr |
| YS051 | h90 | Δmei4::ura4$^{+}$ rec8-S552P-GFP-bsdr Δwpl1::hygr |
| **Figure S5E** | | |
| YS282 | h+/h- | mei4::ura4$^{+}$/mei4::ura4$^{+}$ leu1/leu1 ura4-D18/ura4-D18 ade6-M210/M216 rec8$^{+}$-GFP-bsdr/rec8$^{+}$-GFP-bsdr rec11-3flag-kanr/rec11-3flag-kanr psm1-3HA-hyg.Br/psm1-3HA-hygr |
| YS275 | h+/h- | mei4::ura4$^{+}$/mei4::ura4$^{+}$ leu1/leu1 ura4-D18/ura4-D18 ade6-M210/M216 rec8-S552P-GFP-bsdr/rec8-S552P-GFP-bsdr rec11-3flag-kanr/rec11-3flag-kanr psm1-3HA-hygr/psm1-3HA-hygr |
| **Figure S5F** | | |
| YS705 | h-/h- | ade6-M216/210 leu1/leu1 ura4-D18/ura4-D18 pat1-L95A<<bleo/pat1-L95A<<bleo caf5::bsdr/caf5::bsdr pap1-del/pap1-del pmd1-del/pmd1-del mfs1-del/mfs1-del bfr1-del/bfr1-del dnf2-del/dnf2-del erg5::ura4$^{+}$/erg5::ura4$^{+}$ +/CO2::mat-Pc-natr rec8$^{+}$-3Flag-kanr/rec8$^{+}$-3Flag-kanr psm1-3HA-hygr/psm1-3HA-hygr |
| YS706 | h-/h- | ade6-M216/210 leu1/leu1 ura4-D18/ura4-D18 pat1-L95A<<bleo/pat1-L95A<<bleo caf5::bsdr/caf5::bsdr pap1-del/pap1-del pmd1-del/pmd1-del mfs1-del/mfs1-del bfr1-del/bfr1-del dnf2-del/dnf2-del erg5::ura4$^{+}$/erg5::ura4$^{+}$ +/CO2::mat-Pc-natr rec8-F204S-3Flag-kanr/rec8-F204S-3Flag-kanr psm1-3HA-hygr/psm1-3HA-hygr |
| **Figure S5G** | | |
| YS604 | h90 | Δmei4::ura4$^{+}$ rec11$^{+}$-GFP-kanr Δwpl1::hygr rec8$^{+}$-3Pk-bsdr |
| YS606 | h90 | Δmei4::ura4$^{+}$ ura4$^{+}$ rec11$^{+}$-GFP-kanr Δwpl1::hygr rec8-F204S-3Pk-bsdr |
| **Figure S5H** | | |
| PC345 | h90 | Δmei4::ura4$^{+}$ rec8$^{+}$-GFP-kanr Δwpl1::hygr |
| YS370 | h90 | Δwpl1::hygr Δmei4::ura4$^{+}$ ade6? rec8-F204S-GFP-bsdr |
| YS373 | h90 | Δwpl1::hygr Δmei4::ura4$^{+}$ ade6? rec8-F204T-GFP-bsdr |
| YS378 | h90 | Δwpl1::hygr Δmei4::ura4$^{+}$ ade6? rec8-F204N-GFP-bsdr |
| YS374 | h90 | Δwpl1::hygr Δmei4::ura4$^{+}$ ade6? rec8-F204D-GFP-bsdr |
| YS371 | h90 | Δwpl1::hygr Δmei4::ura4$^{+}$ ade6? rec8-F204A-GFP-bsdr |
| YS379 | h90 | Δwpl1::hygrΔmei4::ura4$^{+}$ ade6? rec8-F204R-GFP-bsdr |
| YS375 | h90 | Δwpl1::hygr Δmei4::ura4$^{+}$ ade6? rec8-F204H-GFP-bsdr |
| YS372 | h90 | Δwpl1::hygr Δmei4::ura4$^{+}$ ade6? rec8-F204C-GFP-bsdr |
| YS377 | h90 | Δwpl1::hygr Δmei4::ura4$^{+}$ ade6? rec8-F204W-GFP-bsdr |
| YS376 | h90 | Δwpl1::hygr Δmei4::ura4$^{+}$ ade6? rec8-F204L-GFP-bsdr |
| **Figure S6A** | | |
| YS547 | h-/h- | ade6-M216/210 leu1/leu1 ura4-D18/ura4-D18 pat1-L95A<<bleo/pat1-L95A<<bleo caf5::bsdr/caf5::bsdr pap1-del/pap1-del pmd1-del/pmd1-del mfs1-del/mfs1-del bfr1-del/bfr1-del dnf2-del/dnf2-del erg5::ura4$^{+}$/erg5::ura4$^{+}$ +/CO2::mat-Pc-natr rec8$^{+}$-3Flag-kanr/rec8$^{+}$-3Flag-kanr |
| YS549 | h-/h- | ade6-M216/210 leu1/leu1 ura4-D18/ura4-D18 pat1-L95A<<bleo/pat1-L95A<<bleo caf5::bsdr/caf5::bsdr pap1-del/pap1-del pmd1-del/pmd1-del mfs1-del/mfs1-del bfr1-del/bfr1-del dnf2-del/dnf2-del erg5::ura4$^{+}$/erg5::ura4$^{+}$ +/CO2::mat-Pc-natr rec8-F204S-3Flag-kanr/rec8-F204S-3Flag-kanr |
| **Figure S6B** | | |

| | | |
|---|---|---|
| YS724 | h-/h- | ade6-M216/210 leu1/leu1 ura4-D18/ura4-D18 pat1-L95A<<bleo/pat1-L95A<<bleo caf5::bsdr/caf5::bsdr pap1-del/pap1-del pmd1-del/pmd1-del mfs1-del/mfs1-del bfr1-del/bfr1-del dnf2-del/dnf2-del erg5::ura4$^{+}$/erg5::ura4$^{+}$ +/CO2::mat-Pc-natr rec8$^{+}$-3Flag-kanr/rec8$^{+}$-3Flag-kanr rec11-3HA-hygr/rec11-3HA-hygr |
| YS728 | h-/h- | ade6-M216/210 leu1/leu1 ura4-D18/ura4-D18 pat1-L95A<<bleo/pat1-L95A<<bleo caf5::bsdr/caf5::bsdr pap1-del/pap1-del pmd1-del/pmd1-del mfs1-del/mfs1-del bfr1-del/bfr1-del dnf2-del/dnf2-del erg5::ura4$^{+}$/erg5::ura4$^{+}$ +/CO2::mat-Pc-natr rec8-F204S-3Flag-kanr/rec8-F204S-3Flag-kanr rec11-3HA-hygr/rec11-3HA-hygr |
| **Figure S7A** | | |
| YS1156 | h90 | rec8$^{+}$-GFP-bsdr rec10$^{+}$-mCherry-natr leu1 (rec11$^{+}$) |
| YS1157 | h90 | rec8$^{+}$-GFP-bsdr rec10$^{+}$-mCherry-natr leu1 rec11-5A ade6-M210 |
| YS1158 | h90 | rec8$^{+}$-GFP-bsdr rec10$^{+}$-mCherry-natr leu1 rec11-5D ade6-M210 |
| YS1162 | h90 | rec8-F204S-GFP-bsdr rec10$^{+}$-mCherry-natr leu1 (rec11$^{+}$) |
| YS1163 | h90 | rec8-F204S-GFP-bsdr rec10$^{+}$-mCherry-natr leu1 rec11-5A ade6-M210 |
| YS1164 | h90 | rec8-F204S-GFP-bsdr rec10$^{+}$-mCherry-natr leu1 rec11-5D ade6-M210 |
| YS1255 | h90 | hygr-Puhp1-3HA-hhp1 Δhhp2::kanr rec10$^{+}$-mCherry-natr rec8$^{+}$-GFP-bsdr |
| YS1258 | h90 | hygr-Puhp1-3HA-hhp1 Δhhp2::kanr rec10$^{+}$-mCherry-natr rec8$^{+}$-GFP-bsdr rec11-5A ade6-M210 |
| YS1261 | h90 | hygr-Puhp1-3HA-hhp1 Δhhp2::kanr rec10$^{+}$-mCherry-natr rec8$^{+}$-GFP-bsdr rec11-5D ade6-M210 |
| **Figure S7B** | | |
| YS1305 | h90 | rec8$^{+}$-3Pk-bsdr ura4 ade6-M26 + pade6-469 TF |
| YS1307 | h90 | rec8-F204S-3Pk-bsdr ura4 ade6-M26 + pade6-469 |
| YS1309 | h90 | Δrec8::kanr ura4 ade6-M26 $^{+}$ pade6-469 |
| **Figure S7C** | | |
| PZ351 | h- | lys3 ura1 |
| JY334 | h+ | leu1 ade6 |
| PK535 | h- | hygr-Puhp1-3HA-hhp1 Δhhp2::kanr ura1 lys3 |
| PK483' | h+ | kanr-Puhp1-3HA-hhp1 Δhhp2::natr leu1 |
| PC398 | h- | rec11-5D ade6-M210 ura1 lys3 |
| PC399 | h+ | rec11-5D ade6-M210 leu1 |
| PC403 | h- | hygr-Puhp1-3HA-hhp1 Δhhp2:kanr rec11-5D ade6-M210 ura1 lys3 |
| PC406 | h+ | hygr-Puhp1-3HA-hhp1 Δhhp2:kanr rec11-5D ade6-M210 leu1 |
| YS414 | h- | ura1 lys3 rec8-F204S-3Pk-bsdr |
| YS420 | h+ | rec8-F204S-3Pk-bsdr leu1 |
| YS1335 | h- | rec8-F204S-3Pk-bsdr ade6 rec11-5D |
| YS1338 | h+ | rec8-F204S-3Pk-bsdr ura1 lys3 leu1 ade6 rec11-5D |
| **Figure S7D** | | |
| YS648 | h90 | rec11$^{+}$-GFP-rec10$^{+}$-bsdr rec25$^{+}$-mCherry-natr Δmei4::hygr rec8$^{+}$-3Pk-kanr Δrec10::bsdr leu1 ade6-M216 |
| YS652 | h90 | rec11$^{+}$-GFP-rec10$^{+}$-bsdr rec25$^{+}$-mCherry-natr Δmei4::hygr rec8-F204S-3Pk-kanr Δrec10::bsdr leu1 ade6-M216 |
| **Supplemental movie1** | | |
| YS686 | h90 | rec8$^{+}$-GFP-bsdr ade6? aur1$^{+}$::htb1-mCherry |
| **Supplemental movie2** | | |
| YS1152 | h90 | rec8$^{+}$-GFP-bsdr ade6? aur1$^{+}$::htb1-mCherry Δwpl1::natr |
| **Supplemental movie3** | | |
| YS688 | h90 | rec8-F204S-GFP-bsdr ade6? aur1$^{+}$::htb1-mCherry |

**Supplementary Table S2.** The mapping statistics of Hi-C data used in this study.

| | wt_VG | | wt_0 h | | wt_2.5 h | |
|---|---|---|---|---|---|---|
| | number | ratio | number | ratio | number | ratio |
| Total read | 28,754,666 | 100% | 16,249,350 | 100% | 15,937,250 | 100% |
| Both aligned | 27,349,934 | 95.1% | 15,531,817 | 95.6% | 15,129,992 | 94.9% |
| Single aligned | 1,040,126 | 3.6% | 566,579 | 3.5% | 645,301 | 4.0% |
| Not aligned | 364,606 | 1.3% | 150,954 | 0.9% | 161,957 | 1.0% |
| no PCR duplicate | 26,700,916 | 92.9% | 15,133,916 | 93.1% | 14,730,402 | 92.4% |
| no repeat | 22,964,751 | 79.9% | 13,064,747 | 80.4% | 12,848,103 | 80.6% |
| enough quality(MapQ>30) | 20,665,646 | 71.9% | 11,769,978 | 72.4% | 11,435,514 | 71.8% |
| inter-chromosome | 2,030,113 | 7.1% | 912,818 | 5.6% | 672,762 | 4.2% |
| >10kb | 10,313,030 | 35.9% | 5,310,495 | 32.7% | 4,863,579 | 30.5% |
| <10kb(same direction) | 2,956,733 | 10.3% | 2,159,416 | 13.3% | 2,036,730 | 12.8% |
| <10kb(+ +) | 1,479,484 | 5.1% | 1,081,442 | 6.7% | 1,019,478 | 6.4% |
| <10kb(- -) | 1,477,249 | 5.1% | 1,077,974 | 6.6% | 1,017,252 | 6.4% |
| <10kb(different direction) | 5,365,532 | 18.7% | 3,387,065 | 20.8% | 3,862,280 | 24.2% |
| <10kb(+ -) | 3,953,954 | 13.8% | 2,446,831 | 15.1% | 3,005,809 | 18.9% |
| <10kb(- +) | 1,411,578 | 4.9% | 940,234 | 5.8% | 856,471 | 5.4% |
| Total usable reads | 15,299,876 | 53.2% | 8,382,729 | 51.6% | 7,573,071 | 47.5% |

| | wpl1_VG | | wpl1_0 h | | wpl1_2.5 h | |
|---|---|---|---|---|---|---|
| | number | ratio | number | ratio | number | ratio |
| Total read | 30,398,124 | 100% | 16,967,822 | 100% | 14,740,364 | 100% |
| Both aligned | 28,697,747 | 94.4% | 16,133,713 | 95.1% | 13,945,509 | 94.6% |
| Single aligned | 1,140,377 | 3.8% | 625,372 | 3.7% | 613,478 | 4.2% |
| Not aligned | 560,000 | 1.8% | 208,737 | 1.2% | 181,377 | 1.2% |
| no PCR duplicate | 27,981,685 | 92.1% | 15,713,563 | 92.6% | 13,580,712 | 92.1% |
| no repeat | 23,624,470 | 77.7% | 13,528,634 | 79.7% | 11,732,848 | 79.6% |
| enough quality(MapQ>30) | 21,286,768 | 70.0% | 12,137,925 | 71.5% | 10,377,414 | 70.4% |
| inter-chromosome | 1,760,436 | 5.8% | 683,852 | 4.0% | 554,525 | 3.8% |
| >10kb | 10,688,635 | 35.2% | 5,721,942 | 33.7% | 4,849,907 | 32.9% |
| <10kb(same direction) | 2,985,521 | 9.8% | 2,138,422 | 12.6% | 1,752,442 | 11.9% |
| <10kb(+ +) | 1,492,420 | 4.9% | 1,069,385 | 6.3% | 877,127 | 6.0% |
| <10kb(- -) | 1,493,101 | 4.9% | 1,069,037 | 6.3% | 875,315 | 5.9% |
| <10kb(different direction) | 5,851,954 | 19.3% | 3,593,582 | 21.2% | 3,220,433 | 21.8% |
| <10kb(+ -) | 4,366,255 | 14.4% | 2,652,440 | 15.6% | 2,422,517 | 16.4% |
| <10kb(- +) | 1,485,699 | 4.9% | 941,142 | 5.5% | 797,916 | 5.4% |
| Total usable reads | 15,434,592 | 50.8% | 8,544,216 | 50.4% | 7,156,874 | 48.6% |

| | rec8_0 h | | rec12_0 h | | rec10_0 h | |
|---|---|---|---|---|---|---|
| | number | ratio | number | ratio | number | ratio |
| Total read | 15,244,266 | 100% | 16,927,207 | 100% | 17,005,389 | 100% |
| Both aligned | 14,543,125 | 95.4% | 16,078,277 | 95.0% | 15,963,985 | 93.9% |

| | | | | | | |
|---|---|---|---|---|---|---|
| Single aligned | 554,650 | 3.6% | 636,979 | 3.8% | 606,402 | 3.6% |
| Not aligned | 146,491 | 1.0% | 211,951 | 1.3% | 435,002 | 2.6% |
| no PCR duplicate | 14,198,767 | 93.1% | 15,653,358 | 92.5% | 15,545,067 | 91.4% |
| no repeat | 12,318,623 | 80.8% | 13,478,558 | 79.6% | 12,997,556 | 76.4% |
| enough quality(MapQ>30) | 11,066,027 | 72.6% | 12,053,142 | 71.2% | 11,835,907 | 69.6% |
| inter-chromosome | 1,178,556 | 7.7% | 898,647 | 5.3% | 812,933 | 4.8% |
| >10kb | 4,732,836 | 31.0% | 5,096,291 | 30.1% | 4,324,649 | 25.4% |
| <10kb(same direction) | 2,058,383 | 13.5% | 2,264,037 | 13.4% | 2,142,074 | 12.6% |
| <10kb(+ +) | 1,029,671 | 6.8% | 1,134,590 | 6.7% | 1,071,010 | 6.3% |
| <10kb(- -) | 1,028,712 | 6.7% | 1,129,447 | 6.7% | 1,071,064 | 6.3% |
| <10kb(different direction) | 3,096,148 | 20.3% | 3,794,009 | 22.4% | 4,556,114 | 26.8% |
| <10kb(+ -) | 2,194,766 | 14.4% | 2,766,416 | 16.3% | 3,639,955 | 21.4% |
| <10kb(- +) | 901,382 | 5.9% | 1,027,593 | 6.1% | 916,159 | 5.4% |
| Total usable reads | 7,969,775 | 52.3% | 8,258,975 | 48.8% | 7,279,656 | 42.8% |

| | **rec8_2.5 h** | | **rec12_2.5 h** | | **rec10_2.5 h** | |
|---|---|---|---|---|---|---|
| | number | ratio | number | ratio | number | ratio |
| Total read | 16,195,762 | 100% | 15,226,513 | 100% | 16,712,894 | 100% |
| Both aligned | 15,371,244 | 94.9% | 14,436,411 | 94.8% | 15,847,437 | 94.8% |
| Single aligned | 660,702 | 4.1% | 598,705 | 3.9% | 674,648 | 4.0% |
| Not aligned | 163,816 | 1.0% | 191,397 | 1.3% | 190,809 | 1.1% |
| no PCR duplicate | 14,964,510 | 92.4% | 14,054,299 | 92.3% | 15,423,905 | 92.3% |
| no repeat | 13,114,533 | 81.0% | 12,265,450 | 80.6% | 13,331,774 | 79.8% |
| enough quality(MapQ>30) | 11,616,964 | 71.7% | 10,904,847 | 71.6% | 11,915,231 | 71.3% |
| inter-chromosome | 1,436,013 | 8.9% | 631,728 | 4.1% | 807,418 | 4.8% |
| >10kb | 4,627,083 | 28.6% | 4,561,265 | 30.0% | 4,818,926 | 28.8% |
| <10kb(same direction) | 2,003,997 | 12.4% | 1,946,183 | 12.8% | 2,083,137 | 12.5% |
| <10kb(+ +) | 1,002,883 | 6.2% | 974,486 | 6.4% | 1,041,809 | 6.2% |
| <10kb(- -) | 1,001,114 | 6.2% | 971,697 | 6.4% | 1,041,328 | 6.2% |
| <10kb(different direction) | 3,549,753 | 21.9% | 3,765,516 | 24.7% | 4,205,592 | 25.2% |
| <10kb(+ -) | 2,711,076 | 16.7% | 2,918,343 | 19.2% | 3,325,364 | 19.9% |
| <10kb(- +) | 838,677 | 5.2% | 847,173 | 5.6% | 880,228 | 5.3% |
| Total usable reads | 8,067,093 | 49.8% | 7,139,176 | 46.9% | 7,709,481 | 46.1% |

| | **rec8-wt_0 h** | | **rec8-F204S_0 h** | |
|---|---|---|---|---|
| | number | ratio | number | ratio |
| Total read | 16,335,306 | 100% | 16,948,118 | 100% |
| Both aligned | 15,577,432 | 95.4% | 16,211,523 | 95.7% |
| Single aligned | 649,453 | 4.0% | 611,228 | 3.6% |
| Not aligned | 108,421 | 0.7% | 125,367 | 0.7% |
| no PCR duplicate | 15,157,154 | 92.8% | 15,777,206 | 93.1% |
| no repeat | 13,273,816 | 81.3% | 13,884,433 | 81.9% |
| enough quality(MapQ>30) | 11,710,163 | 71.7% | 12,467,844 | 73.6% |
| inter-chromosome | 1,076,278 | 6.6% | 1,259,416 | 7.4% |

| | | | | |
|---|---|---|---|---|
| >10kb | 5,406,907 | 33.1% | 5,129,769 | 30.3% |
| <10kb(same direction) | 2,009,999 | 12.3% | 2,227,957 | 13.1% |
| <10kb(+ +) | 1,006,278 | 6.2% | 1,113,764 | 6.6% |
| <10kb(- -) | 1,003,721 | 6.1% | 1,114,193 | 6.6% |
| <10kb(different direction) | 3,216,801 | 19.7% | 3,850,573 | 22.7% |
| <10kb(+ -) | 2,298,012 | 14.1% | 2,885,592 | 17.0% |
| <10kb(- +) | 918,789 | 5.6% | 964,981 | 5.7% |
| Total usable reads | 8,493,184 | 52.0% | 8,617,142 | 50.8% |

| | **rec8-wt_2.5 h** | | **rec8-F204S_2.5 h** | |
|---|---|---|---|---|
| | number | ratio | number | ratio |
| Total read | 15,500,470 | 100% | 15,354,750 | 100% |
| Both aligned | 14,580,292 | 94.1% | 14,482,078 | 94.3% |
| Single aligned | 771,184 | 5.0% | 755,307 | 4.9% |
| Not aligned | 148,994 | 1.0% | 117,365 | 0.8% |
| no PCR duplicate | 14,176,208 | 91.5% | 14,086,944 | 91.7% |
| no repeat | 12,364,454 | 79.8% | 12,337,169 | 80.3% |
| enough quality(MapQ>30) | 10,623,721 | 68.5% | 10,576,924 | 68.9% |
| inter-chromosome | 792,184 | 5.1% | 1,237,957 | 8.1% |
| >10kb | 4,940,006 | 31.9% | 4,759,129 | 31.0% |
| <10kb(same direction) | 1,846,360 | 11.9% | 1,802,762 | 11.7% |
| <10kb(+ +) | 924,506 | 6.0% | 901,558 | 5.9% |
| <10kb(- -) | 921,854 | 5.9% | 901,204 | 5.9% |
| <10kb(different direction) | 3,044,989 | 19.6% | 2,776,949 | 18.1% |
| <10kb(+ -) | 2,235,685 | 14.4% | 1,979,449 | 12.9% |
| <10kb(- +) | 809,304 | 5.2% | 797,500 | 5.2% |
| Total usable reads | 7,578,550 | 48.9% | 7,799,848 | 50.8% |

**Legend of Supplementary Movies**

**Supplementary Movie 1.** The horsetail nuclear movement in *rec8* wild type. Time-lapse observation (5-min intervals) during meiosis from the horsetail stage in wild type with Rec8-GFP. Chromosomes are labeled by ectopic expression of histone H2B-mCherry. The vast majority of Rec8-GFP proteins were destroyed at the onset of anaphase I.

**Supplementary Movie 2.** The horsetail nuclear movement in *wpl1*Δ mutant. Time-lapse observation (5-min intervals) during meiosis from the horsetail stage in *wpl1*Δ mutant with Rec8-GFP. Chromosomes are labeled by ectopic expression of histone H2B-mCherry. The vast majority of Rec8-GFP proteins were destroyed at the onset of anaphase I.

**Supplementary Movie 3.** The horsetail nuclear movement in *rec8-F204S* mutant.
Time-lapse observation (5-min intervals) during meiosis from the horsetail stage in *rec8-F204S* mutant with Rec8-GFP. Chromosomes are labeled by ectopic expression of histone H2B-mCherry. The vast majority of Rec8-GFP proteins were destroyed at the onset of anaphase I.
